# Multiplexed Analysis of the Secretin-like GPCR-RAMP Interactome

**DOI:** 10.1101/597690

**Authors:** Emily Lorenzen, Tea Dodig-Crnković, Ilana B. Kotliar, Elisa Pin, Emilie Ceraudo, Roger D. Vaughan, Mathias Uhlèn, Thomas Huber, Jochen M. Schwenk, Thomas P. Sakmar

## Abstract

Although receptor activity-modifying proteins (RAMPs) have been shown to modulate the functions of several different G protein-coupled receptors (GPCRs), potential direct interactions among the three known RAMPs and hundreds of GPCRs has never been investigated. We engineered three epitope-tagged RAMPs and 23 epitope-tagged GPCRs, focusing on the secretin-like family of GPCRs, and developed a suspension bead array (SBA) immunoassay designed to detect RAMP-GPCR complexes. We then used 64 antibodies raised against native RAMPs and GPCRs, along with four antibodies targeting the epitope tags, to multiplex the SBA assay to detect and measure all possible combinations of interaction among the 23 GPCRs and three RAMPs. The results of the SBA assay provide a complete interactome of secretin-like GPCRs with RAMPs. We demonstrate direct interaction of previously reported secretin-like GPCRs whose functions are modulated by RAMPs. We also discovered novel sets of GPCR-RAMP interacting pairs, and found additional secretin-like GPCRs, chemokine receptors and orphan receptors that interact with RAMPs. Using *in situ* proximity ligation assay, we verified a subset of these novel GPCR-RAMP interactions in cell membranes. In total, we found GPCR-RAMP interactions for the majority of the 23 GPCRs tested. Each GPCR interacted with either all three RAMPs or with RAMP2 and RAMP3, with the exception of one GPCR that interacted with just RAMP3. In summary, we describe an SBA strategy that will be useful to search for GPCR-RAMP interactions in cell lines and tissues, and conclude that GPCR-RAMP interactions are more common than previously appreciated.

## Introduction

G protein-coupled receptors (GPCRs) are a large family of cell-surface receptors that are commonly targeted for drug development. Emerging evidence suggests that the surface expression and activity of GPCRs can be modulated by receptor activity-modifying proteins (RAMPs), a ubiquitously-expressed family of single-pass membrane proteins with only three members. First discovered as a necessary component for the function of calcitonin-like receptor (CALCRL), RAMPs were shown to alter ligand specificity of both CALCRL and calcitonin receptor (CALCR) by dictating to which ligand they respond, *(1–3)*. Interaction of RAMPs with GPCRs has since been demonstrated with several additional members of the secretin-like GPCRs. RAMPs can influence several features of GPCR biology, including trafficking to the cell membrane, ligand specificity, downstream signaling and recycling *(4–11)*. However, many of the secretin-like GPCR-RAMP interactions remain unverified, and most of the interactions have not been demonstrated using a direct binding assay. In addition, prior reports show conflicting results regarding whether certain secretin-like GPCRs form complexes with RAMPs.

There are also suggestions that RAMPs form complexes with a broader set of GPCRs. A single GPCR from the glutamate-like family, calcium-sensing receptor (CaSR), and a single GPCR from the rhodopsin-like family, G protein-coupled estrogen receptor 1 (GPER1), have been reported to interact with at least one RAMP *(4–6)*. Several lines of evidence from disease-associated pathologies, tissue expression and knock out mice indicate that the breadth of GPCR-RAMP interactions is not yet fully appreciated *(12, 13)*. Furthermore, we demonstrated recently that GPCRs and RAMPs globally co-evolved and display correlated mRNA levels across human tissues, which suggests the likelihood for more widespread GPCR-RAMP interactions *(14)*. We also showed concordance between mRNA levels for a subset of GPCRs and RAMP2 using a single-cell MERFISH (multiplexed error reduced fluorescence in situ hybridization) method *(15)*.

Given that there are ∼400 non-olfactory GPCRs that might in principle interact with each of the three RAMPs, the ideal experimental system should be high-throughput and provide quantitative assessment of GPCR-RAMP complexes. Towards this end, we employed a suspension bead array (SBA) approach. The SBA strategy is based on the use of magnetic microspheres with different bar-codes. Each bar-coded bead population is coupled to a specific antibody (Ab), allowing the parallel capture and detection of multiple unique, identifiable protein epitopes from a complex mixture of proteins in solution. The SBA technology in combination with the Human Protein Atlas (HPA) Ab collection has been successfully implemented to identify soluble proteins in human serum samples *(16, 17)*. Here we demonstrate an experimental approach to define the secretin-like GPCR-RAMP interactome, and develop an assay capable of testing the hypothesis that GPCRs and RAMPs interact on a global level.

## Results

### Preparation of SBA and lysates to screen for GPCR-RAMP complexes

To adapt the SBA method and develop an assay to measure GPCR-RAMP interactions, we focused on the family of secretin-like GPCRs, and eight additional GPCRs, to screen for their ability to interact with each of the three RAMPs. All of the secretin-like receptors were included in the test set because this GPCR family encompasses the majority of GPCR-RAMP interactions described to date. Based on an earlier co-evolution and co-expression analyses we also included GPR4 and GPR182 from the rhodopsin-like GPCR family, and ADGRF5 from the adhesion GPCR family *(14)*. In addition, the chemokine cluster of GPCRs demonstrated significantly higher median co-expression with RAMP2 and RAMP3 in comparison with all GPCRs (fig. S1A). Thus, we included members of the chemokine receptor family: C-C chemokine receptor 5 (CCR5), C-C chemokine receptor 7 (CCR7), C-X-C chemokine receptor 3 (CXCR3), C-X-C chemokine receptor 4 (CXCR4), and atypical chemokine receptor 3 (ACKR3), also known as C-X-C chemokine receptor 7 (CXCR7). A list of each of the 23 GPCRs chosen for SBA analysis is presented in Table 1 and a graphical representation of their placement on the GPCR phylogenetic tree is presented in fig. S1B.

**Table 1.** List of 23 GPCRs included in this study. Listed are the common abbreviations of each GPCR, GPCR family designation, and engineered epitope tag on the N- and C-terminal tails. GPCRs are listed in alphabetical order. Secretin-family receptors are shown in bold. Y, yes; N, No.

| GPCR | Abbreviation | GPCR family | N-terminal HA epitope tag | C-terminal 1D4 epitope tag | Previously Published RAMP interaction |
| --- | --- | --- | --- | --- | --- |
| Atypical chemokine receptor 3/C-X-C chemokine receptor 7 | ACKR3/<br>CXCR7 | Rhodopsin | Y | Y | Unknown |
| **Pituitary adenylate cyclase-activating polypeptide type 1** | ADCYAP1R1 | Secretin | Y | Y | Unknown |
| Adhesion G protein-coupled receptor F5 | ADGRF5 | Adhesion | Y | Y | Unknown |
| **Calcitonin receptor-like receptor** | CALCRL | Secretin | Y | Y | RAMP1, 2 and 3(1) |
| **Calcitonin receptor** | CALCR | Secretin | Y | Y | RAMP1, 2 and 3 (2, 3) |
| C-C chemokine receptor type 5 | CCR5 | Rhodopsin | N | Y | Unknown |
| C-C chemokine receptor type 7 | CCR7 | Rhodopsin | N | Y | Unknown |
| **Corticotropin-releasing hormone receptor 1** | CRHR1 | Secretin | Y | Y | RAMP2 (40) |
| **Corticotropin-releasing hormone receptor 2** | CRHR2 | Secretin | Y | Y | Unknown |
| C-X-C chemokine receptor type 3 | CXCR3 | Rhodopsin | Y | N | Unknown |
| C-X-C chemokine receptor type 4 | CXCR4 | Rhodopsin | N | Y | Unknown |
| **Gastric inhibitory polypeptide receptor** | GIPR | Secretin | Y | Y | Unknown |
| **Glucagon receptor** | GCGR | Secretin | Y | Y | RAMP2 (8, 9, 11) |
| **Growth hormone releasing hormone** | GHRHR | Secretin | Y | Y | None (9) |
| **Glucagon-like receptor 1** | GLP1R | Secretin | Y | Y | None (8–10) |
| **Glucagon-like receptor 2** | GLP2R | Secretin | Y | Y | None (9) |
| G protein-coupled receptor 4 | GPR4 | Rhodopsin | Y | N | Unknown |
| G protein-coupled receptor 182 | GPR182 | Rhodopsin | Y | N | Unknown |
| **Parathyroid hormone receptor 1** | PTH1R | Secretin | Y | Y | RAMP2 (9) |
| **Parathyroid hormone receptor 2** | PTH2R | Secretin | Y | Y | RAMP3 (9) |
| **Secretin receptor** | SCTR | Secretin | Y | Y | RAMP3 (7) |
| **VIP and PACAP receptor 1** | VIPR1 | Secretin | Y | Y | RAMP1, 2 and 3(9) |
| **VIP and PACAP receptor 2** | VIPR2 | Secretin | Y | Y | RAMP1, 2, and 3 (10) |

Our aim was to create an SBA comprising a set of both GPCR-specific and RAMP-specific Abs, as well as monoclonal antibodies (mAbs) against epitope tags that were engineered onto expressed GPCR and RAMP constructs. The complete list of all anti-GPCR and anti-RAMP Abs coupled to the beads is presented in table S1. A total of 55 Abs from the HPA were available to target 21 of the 23 GPCRs in the test set. The HPA Abs were produced by using 50 to 150 amino-acid-long peptides as immunogens to raise rabbit polyclonal Abs *(18)*. The nine anti-RAMP Abs employed were obtained from the HPA and various commercial sources. In addition, beads were coupled with four, validated anti-epitope tag mAbs (anti-HA, anti-1D4, anti-FLAG and anti-OLLAS) *(19, 20)*. These four beads were included to capture GPCRs or RAMPs that were engineered to contain an epitope-fusion tag (Fig. 1A). Each of the 23 GPCRs was epitope-tagged at its N-terminal tail with HA epitope and/or at its C-terminal tail with 1D4 epitope (Table 1). Each of the three RAMPs was tagged with FLAG epitope and OLLAS epitope at its N- and C-terminal tail, respectively. For controls, we used rabbit IgG, mouse IgG and uncoupled beads. The final SBA comprised 70 unique Abs coupled to unique bead IDs, and pooling created the basis for the multiplex assay (Fig. 1B).

**Fig. 1.**
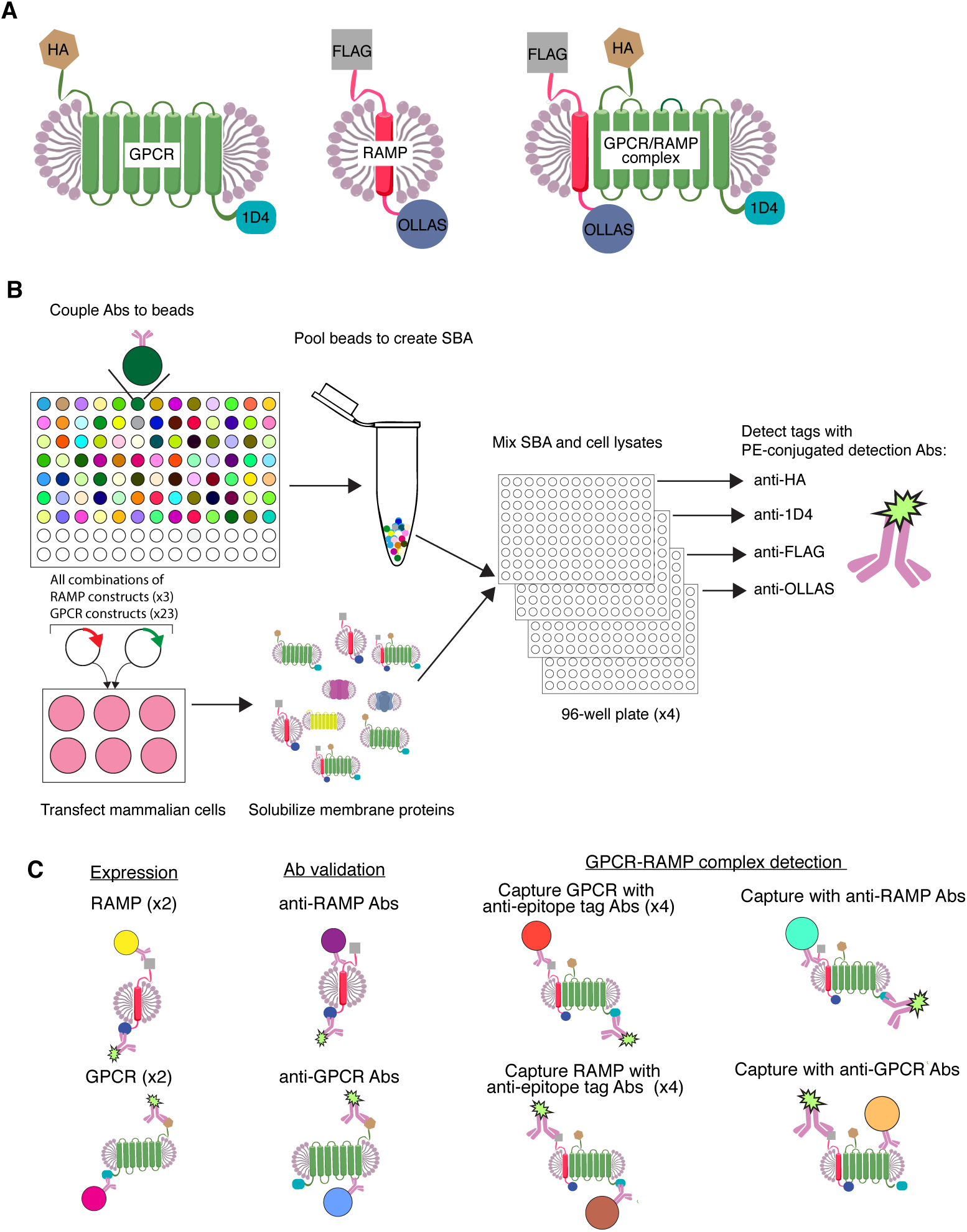
Schematic of SBA assay procedure. **(A)** The GPCRs were epitope-fusion tagged at their N-terminal and C-terminal tails with HA and/or 1D4, respectively. The three RAMPs were tagged at their N-terminal and C-terminal tails with FLAG and OLLAS, respectively. **(B)** Unique Abs were coupled to different bar-coded beads to create an SBA with 70 different populations of capture beads and the beads were subsequently pooled. DNA constructs encoding each of three epitope-tagged RAMPs and 23 epitope-tagged GPCRs were co-transfected in HEK293F cells such that all possible combinations of GRCPs and RAMPs were represented. The cells were solubilized in a detergent solution, which resulted in heterogeneous mixtures of solubilized proteins including the RAMPs, GPCRs and putative GPCR-RAMP complexes. An aliquot of each cell lysate was incubated with an aliquot of SBA. Four identical assay plates were prepared in this manner. Following wash steps, a different PE-conjugated anti-epitope tag detection mAb was added to each of the four plates. **(C)** A Luminex 3D FlexMap instrument was used to measure the reporter fluorescence produced by the PE-conjugated detection mAb while simultaneously reading the bar-code of each individual bead. From a single well, the specificity of RAMP Abs and GPCR Abs could be determined. Simultaneously, GPCR-RAMP complexes could be detected using either anti-epitope tag Abs, anti-GPCR Abs or anti-RAMP Abs.

Each of the three epitope-tagged RAMP constructs was expressed in HEK293F cells alone or co-expressed with each of the 23 epitope-tagged GPCR constructs. Following transfection, each cell preparation was solubilized in a dodecyl maltoside detergent solution in order to create a heterogeneous micelle mixture of RAMPs, GPCRs and GPCR-RAMP complexes, in addition to other cellular protein components (Fig. 1B). This strategy created a set of 73 unique lysates co-transfected with each combination of the RAMPs and 23 GPCRs, plus three RAMP controls, and empty-vector, or mock, transfected controls.

Next, we distributed each lysate in duplicate into 96-well plates and added aliquots of the pooled SBA to each well. Four replicate plates were processed in parallel, one for each detection Ab. For detection of the proteins or protein complexes bound to the beads, each plate was then incubated with a different anti-epitope tag detection Ab (a PE-conjugated version of either anti-HA, anti-FLAG, anti-1D4 or anti-OLLAS) (Fig. 1B). The fluorescence from the PE-conjugated detection Abs was measured by a flow cytometer (Luminex FlexMap 3D) and matched to the bar-code of each individual bead. The SBA strategy allowed us to evaluate protein expression, Ab specificity and GPCR-RAMP complex formation in a single experiment (Fig. 1C).

We first determined that the epitope-tagged GPCRs and RAMPs could be captured and detected by mAbs targeting the epitope tags. Expression of each dual-epitope tagged RAMP was demonstrated by capture with one of the anti-epitope tag mAb beads (OLLAS or FLAG) and detected with the PE-conjugated version of the other anti-epitope tag mAb (OLLAS or FLAG) (fig. S2A and B). Similarly, expression of each dual-epitope tagged GPCR was demonstrated by capture and subsequent detection with the anti-1D4 and anti-HA mAbs (fig. S2C and D).

#### Capture and detection of GPCR-RAMP complexes using anti-epitope tag mAbs

Having validated that both GPCRs and RAMPs can be captured with one of the anti-epitope tag beads and detected with one of four PE-conjugated, anti-epitope tag mAbs, it follows that we should also be able to detect GPCR-RAMP complexes that were captured by the Abs of the SBA. In the basic experimental setup using epitope tag mAbs, there are eight possible capture-detection mAb pairs that can be used to capture or detect either a RAMP or a GPCR in a given GPCR-RAMP complex: 1D4-FLAG, 1D4-OLLAS, HA-FLAG, HA-OLLAS, FLAG-1D4, FLAG-HA, OLLAS-1D4 and OLLAS-HA (Fig. 2A to H). For single epitope-tagged GPCRs, four capture-detection mAb pairs could be used to detect a GPCR-RAMP complex.

**Fig. 2.**
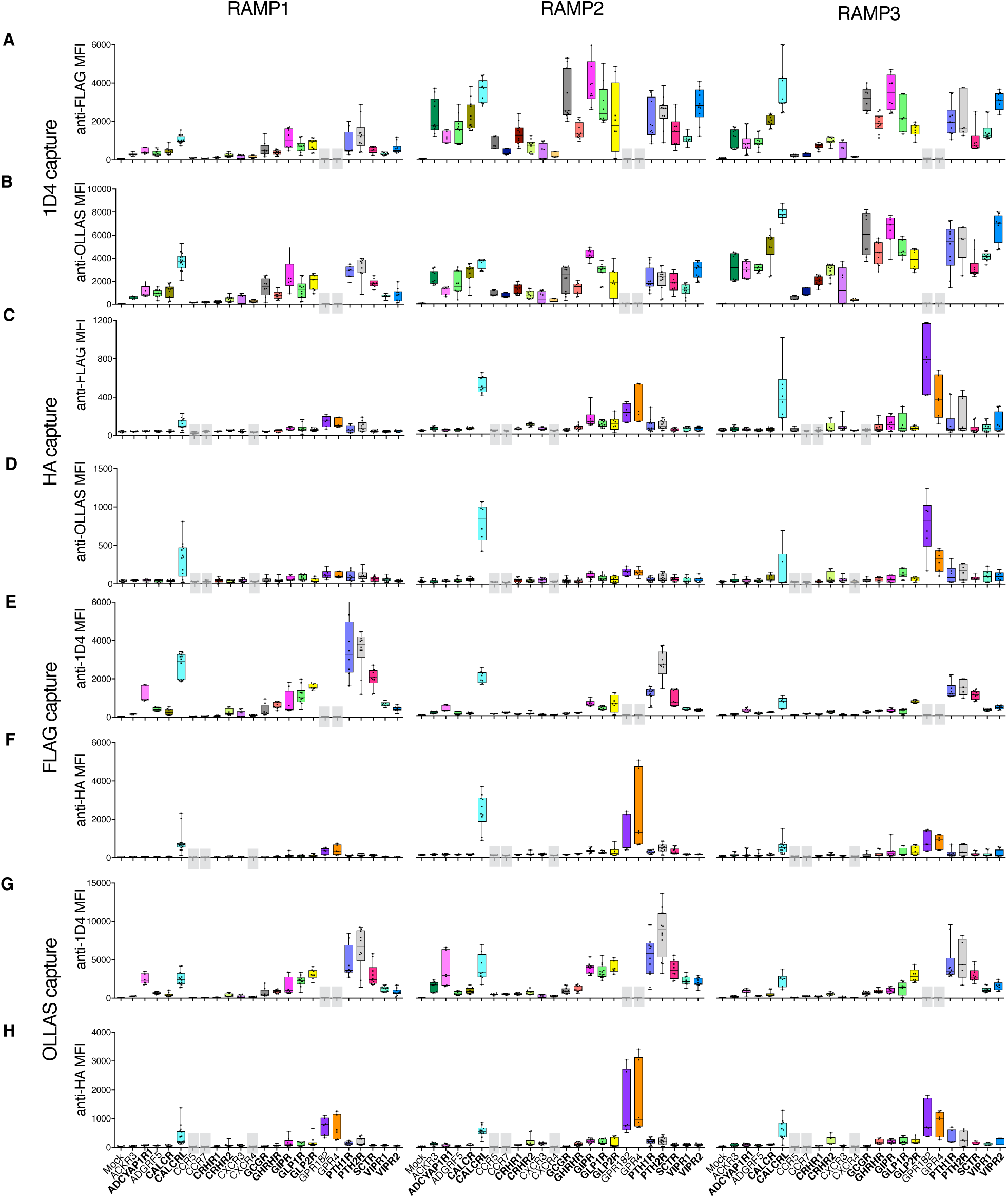
Capture and detection of GPCR-RAMP complexes using anti-epitope tag mAbs. Lysates from cells transfected with each epitope-tagged RAMP construct and co-transfected with each epitope-tagged GPCR construct were incubated with the SBA, which included beads that were conjugated to mAbs targeting the four tags. Complexes were captured in multiplex fashion using one of the four mAbs. There are eight possible capture-detection schemes. The GPCR is captured using anti-1D4 mAb and the GPCR-RAMP complex is detected using **(A)** PE-conjugated FLAG mAb or **(B)** PE-conjugated anti-OLLAS mAb. The GPCR is also captured using anti-HA mAb and the GPCR-RAMP complex is detected using **(C)** PE-conjugated anti-FLAG mAb or **(D)** PE-conjugated anti-OLLAS mAb. The RAMP is captured using anti-FLAG mAb and the GPCR-RAMP complex is detected using **(E)** PE-conjugated 1D4 mAb or **(F)** PE-conjugated anti-HA mAb. The RAMP is captured using anti-OLLAS mAb and the GPCR-RAMP complex is detected using **(G)** PE-conjugated 1D4 mAb or **(H)** PE-conjugated anti-HA mAb. Data are presented for each of the three RAMPs. GPCR names are listed at the bottom of each RAMP panel and the boxes are color coded. The occasional grey box indicates that the GPCR did not have the appropriate epitope tag to be captured or detected. The labels in bold correspond to secretin-like GPCRs. Data are median fluorescence intensity (MFI) and represent measurements obtained from at least three experiments performed in duplicate. The box plots represent the maximum and minimum extents of the measured values and all data points are graphed.

Several GPCRs formed complexes with one or more of the RAMPs, including CALCRL. CALCRL has previously been shown to form a stable complex which each RAMP *(21)*. The significance of the detection signal for the complex depended on the capture-detection Ab pair used (table S2). We calculated an overall statistic for the significance of each GPCR-RAMP complex pair (table S3).

#### Validation of Abs targeting RAMPs and GPCRs

Our next goal was to validate the SBA approach as a method to identify GPCR-RAMP interactions without having to rely upon the use of epitope tags engineered into both the GPCR and the RAMP. Thus, we needed to assess the available protein-specific Abs to each GPCR and each RAMP, which were obtained from either commercial sources or HPA. In order to validate the utility of each anti-RAMP Ab, the cell lysates containing epitope-tagged RAMPs were incubated with the SBA, which included nine anti-RAMP Abs. The captured RAMPs were detected using the PE-conjugated OLLAS and PE-conjugated FLAG mAbs. We found that five out of the nine HPA and commercial anti-RAMP Abs (55%) selectively captured their intended RAMP. In addition, each anti-RAMP Ab showed little cross-reactivity with the other two non-targeted RAMPs (fig. S3). Validated Abs are underlined in table S1.

In parallel, the collection of anti-GPCR Abs was validated for GPCR capture using cell lysates containing the 23 epitope-tagged GPCRs. In this case, the detection Abs used were anti-1D4 and anti-HA. When using either PE-conjugated anti-1D4 or PE-conjugated anti-HA as the detection Ab, 31 out of the 55 anti-GPCR Abs (56%) tested captured their intended GPCR targets (Fig. 3). Overall, we found that the anti-HA mAb was not as sensitive to detect GPCRs as the anti-1D4 detection mAb. To judge the ability of an Ab to capture its intended GPCR target, we used a Z-score cutoff of 1.645 (corresponding to the 95% confidence interval of a single-tailed Z-test). We found that at least one HPA Ab for 19 of the 21 GPCRs (90%) included in this study, and with HPA Abs available, fulfilled the criteria (table S1). We also determined the potential cross-reactivity of all anti-GPCR Abs with unintended GPCR targets and found that none of the validated Abs demonstrated cross-reactivity (fig. S4).

**Fig. 3.**
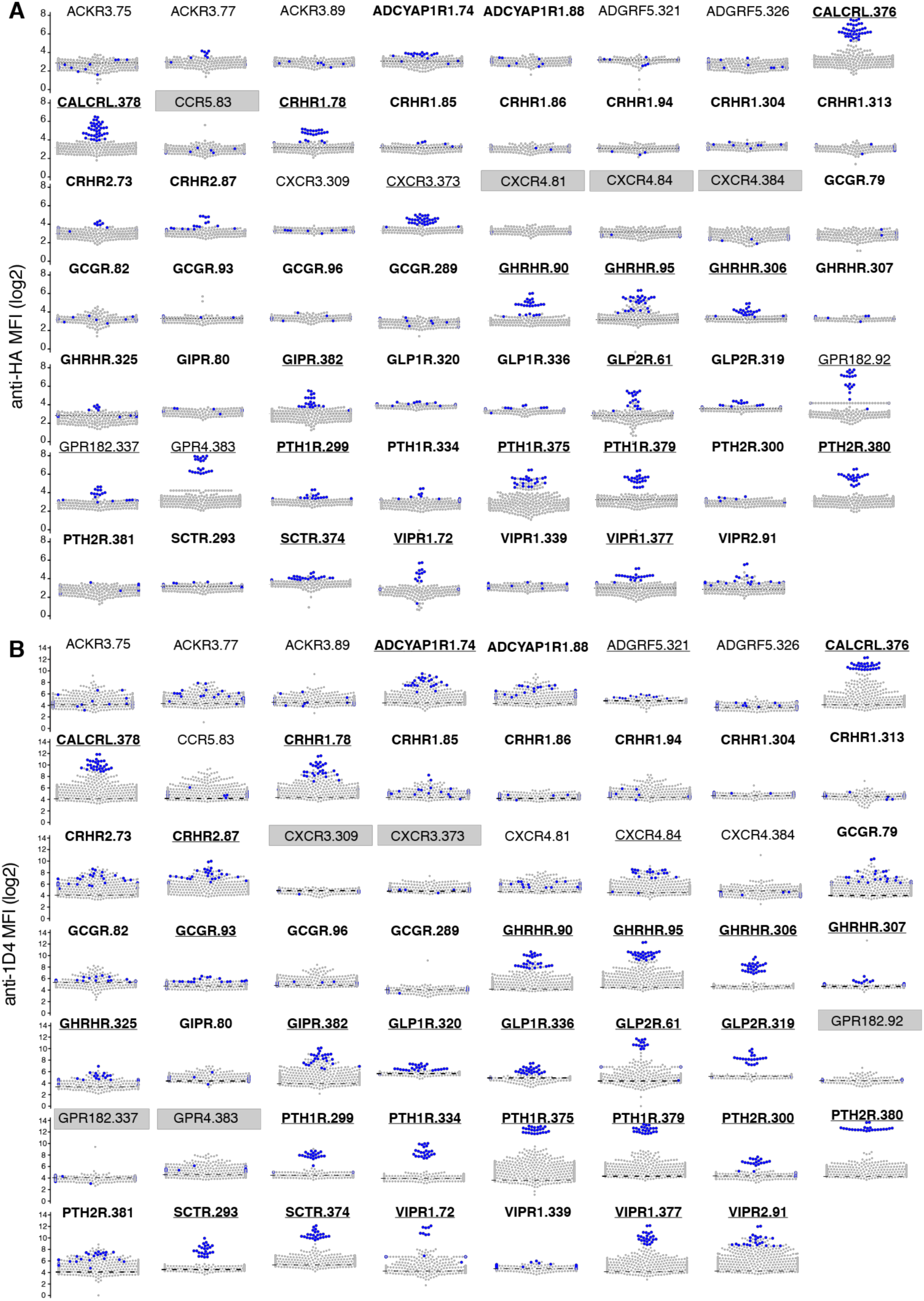
Validation of Abs used to capture GPCRs. In order to validate anti-GPCR Abs, lysates from cells transfected with each epitope-tagged GPCR construct (HA and/or 1D4) were incubated with the SBA, which included beads conjugated with 55 Abs targeting 21 GPCRs. **(A)** PE-conjugated anti-HA and **(B)** PE-conjugated anti-1D4 were used to detect any GPCRs captured by the beads. GPCRs are shown in alphabetical order and the labels in bold correspond to secretin-like GPCRs. Each dot in each bee-swam plot represents a data point from one experiment. The blue dots indicate signal from lysates containing the intended GPCR target, while grey dots indicate signal from lysates containing an of the other epitope-tagged GPCR targets. Data are median fluorescence intensity (MFI) and representative of at least 200 experiments, each performed in duplicate. The occasional grey box indicates that the GPCR did not have the appropriate epitope tag to be captured or detected. At a statistical significance of p*≤*0.05 we validated a total of 31 capture Abs, with at least one capture Ab for 19 of the 21 GPCRs studied. Validated Abs are underlined. Bead ID numbers are listed after each GPCR name and the corresponding Ab name is provided in table S1.

#### GPCR-RAMP complexes captured by validated anti-GPCR and anti-RAMP Abs

Next, we explored the broader utility of using the anti-GPCR and anti-RAMP Abs to capture directly the GPCR-RAMP complexes. We examined the beads that were coupled to vanti-GPCR Abs and measured signals arising from a bound RAMP using both PE-conjugated anti-FLAG and PE-conjugated anti-OLLAS mAbs (fig. S5). In this experimental design, GPCRs can be captured through their native sequence, ablating the need to create epitope-tagged constructs. With few exceptions (<0.05%) the majority of signals associated with complex capture using the anti-GPCR Abs that reached a Z-score *≥* 1.645 were obtained from lysates containing the target GPCR.

We next determined whether the data obtained using anti-GPCR capture Abs recapitulated the results obtained with the epitope-tag capture methods (table S3). Overall, the statistical analysis shows that the results obtained using anti-GPCR Abs to capture the GPCR-RAMP complex corroborates the results obtained when using epitope tags for capture (fig. S6). The population of false negatives arising from the anti-GPCR Ab data is very small. The anti-GPCR Abs of the SBA were also able to capture the majority of the GPCR-RAMP complexes as demonstrated when using an OLLAS detection Ab and with validated anti-GPCR Abs (fig. S6D).

#### Detection of GPCR-RAMP complexes using an SBA assay

We used several detection and capture Ab pairs in a multiplexed fashion to identify GPCR-RAMP complexes. The complexes were captured through epitope tags engineered onto the GPCR or RAMP, and PE-conjugated anti-epitope tag mAbs were used to detect the putative interacting partner. In this experimental setup there were eight combinations of capture/detection pairs to identify complexes of dual-epitope tagged GPCRs with RAMPs, or four capture/detection pairs to identify complexes of single-epitope-tagged constructs. In addition, many of the Abs validated to capture the GPCRs or RAMPs also captured the GPCR-RAMP complexes. These data were collected in multiplexed fashion. A summary of the results obtained from all of the combinations of capture/detection Ab pairs employed are presented graphically in Fig. 4. Generally, the GPCRs that exhibit complex formation with RAMPs either form complexes with all three RAMPs, or RAMP2 and RAMP3. Only one GPCR demonstrated complex formation with just one RAMP, CRHR2 with RAMP3.

**Fig. 4.**
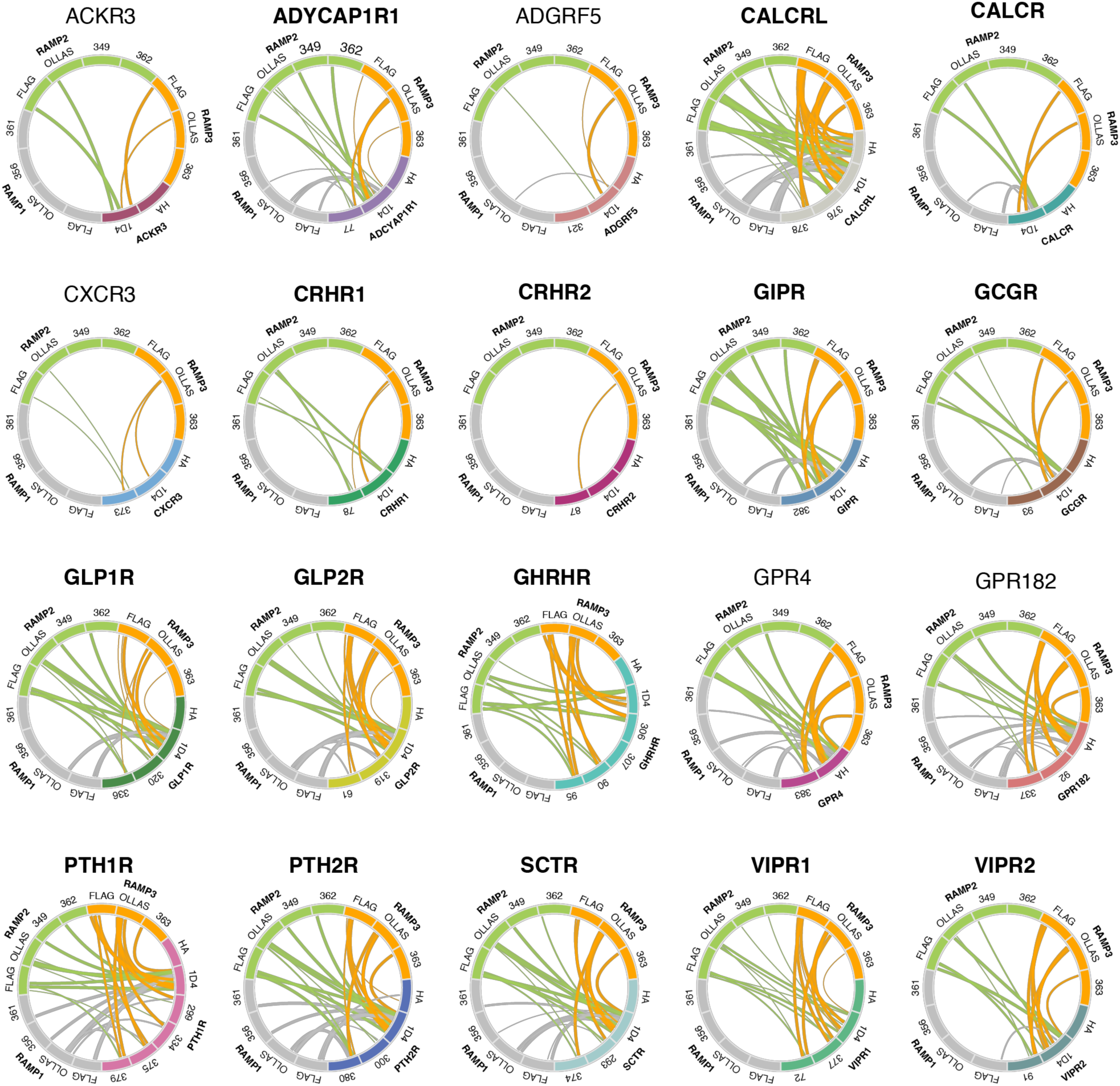
Graphical summary of GPCR-RAMP complexes detected using an SBA assay. Each circle depicts a unique GPCR along with each of the three RAMPs. Each GPCR is labeled and color coded. RAMP1 is colored grey, RAMP2 is colored lime, RAMP3 is colored tangerine. Curved lines within the circles show GPCR-RAMP interactions, and the thicknesses of the lines shows relative statistical significance (see below). The small labels around the circumference indicate the antibodies used for the SBA experiments. In total four anti-epitope tag Abs, 31 validated Abs to the 23 GPCRs included in the study, and five validated Abs to the three RAMPs are shown. Three GPCRs tested (CCR5, CCR7 and CXCR4) did not form complexes with RAMPs and are not shown here. The statistical significance derived for the particular interaction using the indicated capture/detection pair is represented by the thickness of the curved lines. P values of p *≤* 0.05 are given an arbitrary thickness of 1, p *≤* 0.01 a thickness of 2, p *≤* 0.001 a thickness of 3, and p *≤* 0.0001 a thickness of 4. Bead ID numbers are listed with each GPCR name and the corresponding validated Ab names are provided in table S1.

#### Validation of a subset of GPCR-RAMP interactions by proximity ligation assay (PLA)

In order to show the presence of selected GPCR-RAMP complexes in membranes, we employed PLA. The PLA method enables detection of protein interactions, which are visualized as fluorescent puncta *(22)*. In principle, each PLA punctum corresponds to a single protein-protein complex that reacts with two primary Abs, one per protein. We employed multiple controls to verify that GPCR-RAMP interactions could be detected by PLA. The CALCRL-RAMP2 complex was used as a positive control because the complex was identified using the SBA assay and has been shown to exist in cell surface membranes in a variety of studies *(1, 2, 21)*. We employed anti-HA and anti-FLAG Abs to detect the extracellular epitopes of CALCRL and RAMP2, respectively. Omitting each of the primary Abs during PLA processing of CALCRL+RAMP2 co-transfected cells measures primary Ab nonspecific binding, while omitting all primary Abs measures nonspecific binding of the PLA probes. Mock transfected cells treated with both Abs served as a negative control.

We compared PLA puncta count from cells that were co-transfected with CALCRL and RAMP2. We observed a significant number of puncta under the PLA conditions when both CALCRL and RAMP2 were co-expressed (Fig. 5A, fig. S7A and B). We verified that the puncta were from plasma membrane interaction and not an artifact of misfolded protein accumulation in the ER by employing a formaldehyde (FA)-based fixation method (fig. S7C). We next used PLA to test GPCR-RAMP2 interactions in cell membranes for a variety of other GPCRs that were also studied using SBA. We focused on quantifying interactions between RAMP2 and CALCRL, PTH1R, GPR182, GPR4 and CXCR3. These GPCRs showed a range of capabilities to interact with RAMP2 in the SBA under conditions of detergent solubilization. The PLA results indicated that there were differences in puncta count among the receptor-RAMP2 pairs studied (Fig. 5B and C). Overall, the results of the PLA on a limited number of GPCR-RAMP pairs were consistent with the results of the SBA.

**Fig. 5.**
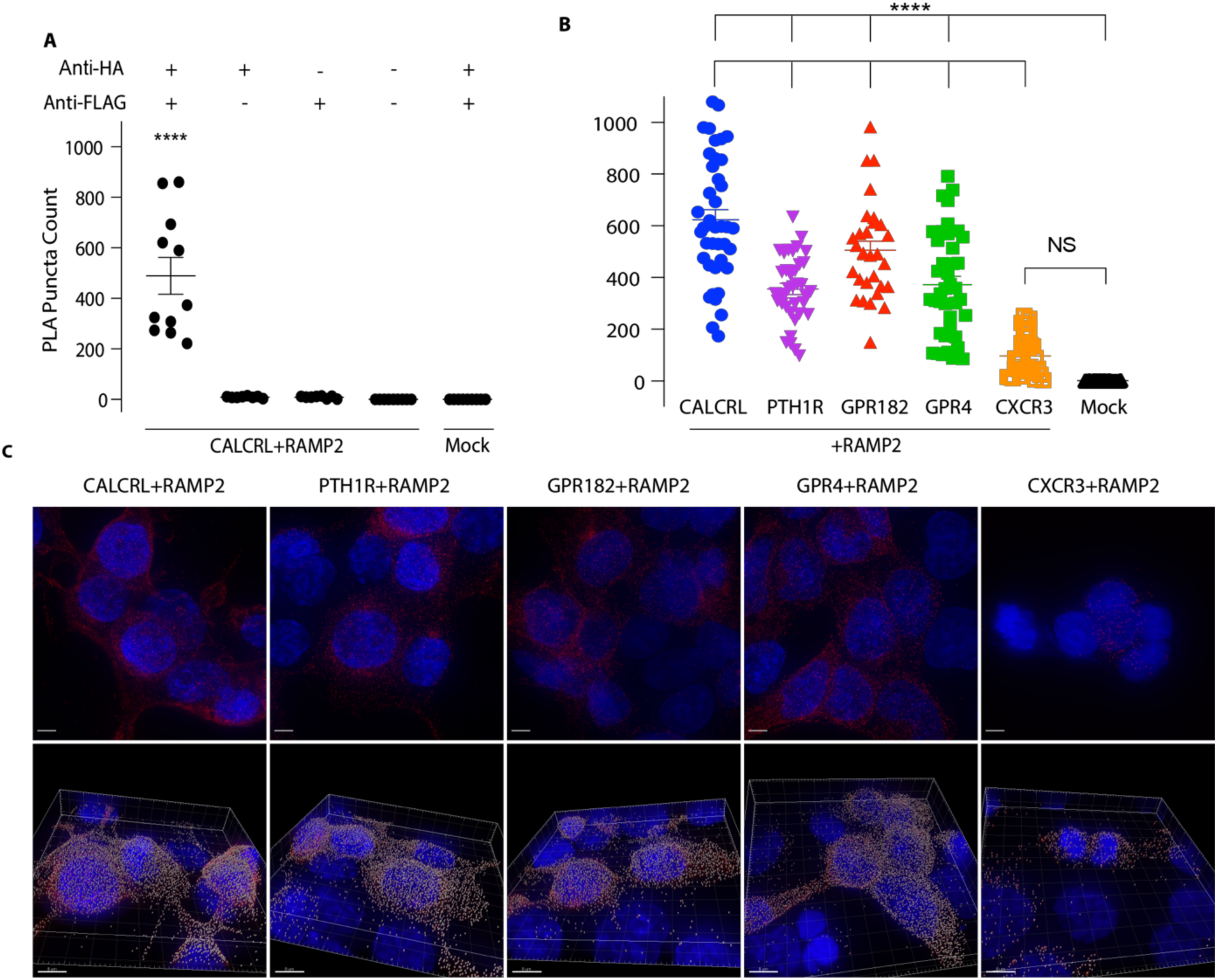
Validation of RAMP-GPCR complex formation in cell membranes using PLA. Cells were co-transfected with epitope-tagged GPCR and RAMP2 then incubated with *α*-HA and *α*-FLAG Abs. PLA was then carried out to quantitate GPCR-RAMP2 interactions. The number of PLA puncta per cell for each Z-stack captured was measured. Each Z-stack is of a different field of view. **(A)** Quantitation of control PLA experiments performed on CALCRL+RAMP2 co-transfected cells. PLA puncta counts were compared between samples that received both primary Abs and samples that received only anti-HA Ab, only anti-FLAG Ab, no primary Abs, or mock transfection with both primary Abs. Data are from two experiments performed with at least three replicates. **(B)** PLA puncta counts for cells co-transfected with RAMP2 and selected GPCRs. Data are from at least three experiments performed with at least five replicates. **(C)** Representative images of cells co-transfected with RAMP2 and selected GPCRs and subjected to PLA. Top row shows maximum projection of Z-stack, which is the maximum signal intensity for each channel at each point across all slices in the Z-stack. The bottom row shows snapshots from puncta quantification performed in Imaris. Scale bars, 5μm top row, 8μm bottom row. Blue = DAPI, red = PLA puncta, grey = Imaris spots. The statistical test for significance used was a one-way ANOVA followed by Dunnett’s multiple comparisons test (****P<0.0001, NS = not significant).

## Discussion

We present a novel method to detect interactions between GPCRs and RAMPs that may have general utility to detect interacting partners of membrane protein more broadly. The method relies on the preparation and validation of Abs for a suspension bead-based assay, which allows multiplexed immunocapture and detection of a large number of discrete proteins from a lysate mixture. The SBA we developed contains uniquely color-coded beads conjugated to four mAbs against epitope tags engineered at the N- and C-terminal tails of the expressed GPCRs and RAMPs, nine Abs against three different RAMPs, 55 Abs against 21 of the 23 GPCRs studied, and control beads. Captured GPCRs, RAMPs, or GPCR-RAMP complexes were detected by mAbs against the engineered epitope tags. In a single multiplexed experiment, we could use a variety of capture and detection strategies to identify complexes. The complexes were captured using the epitope tags on the GPCRs and detected using the epitope tags on the RAMPs, and *vice versa*. Finally, anti-GPCR and anti-RAMP Abs complexes were captured by protein-specific Abs, demonstrating the possibility to detect these complexes without the need for protein engineering to introduce the epitope tags. We found that the results obtained using validated anti-GPCR or anti-RAMP Abs were concordant to those obtained using the anti-epitope tag mAbs for complex capture.

Using the SBA assay strategy, we identified previously reported secretin-like GPCR-RAMP complexes. For example, we show that CALCRL forms stable complexes with each of the three RAMPs, which have been well characterized in the literature *(21)*. In addition, we also detected and confirmed the formation of other complexes between secretin-like GPCRs and RAMPs that have been previously reported: CALCR with all three RAMPs, CRHR1 with RAMP2, GCGR with RAMP2, PTH1R with RAMP2, PTH2R with RAMP3, SCTR with RAMP3, VIPR1 with RAMP2 and RAMP3, and VIPR2 with all three RAMPs *(2, 3, 7–11)*. However, we failed to observe one previously reported complex, VIPR1 with RAMP1. The presence of a putative VIPR1-RAMP1 complex was inferred earlier because the cell surface expression of RAMP1 increased upon co-expression of VIPR1 *(9)*. The majority of the GPCR-RAMP interactions reported earlier were also based upon reciprocal effects of heterologous over-expression, and to our current knowledge, not on any type of direct binding assay. Many of the GPCR-RAMP interactions had remained unverified by other experimental methods. Our findings using the SBA assay system to capture and detect actual GPCR-RAMP complexes appear to validate most of the earlier indirect findings. The results also validate the Abs and assay procedure as a robust method to detect and quantitate the presence of GPCR-RAMP complexes from cell lysates.

In addition, we discovered several new secretin-like GPCR-RAMP complexes that have not been described previously. We found that GIPR and ADCYAP1R1 formed complexes with all three RAMPs. In contrast, CRHR2 showed weak complex formation with RAMP3, but not with RAMP1 and RAMP2. To our knowledge, GIPR, ADCYAP1R1 and CRHR2 have not been studied earlier with respect to their ability to interact with RAMPs. In addition, we found that several secretin-like GPCRs that were previously reported to not interact with RAMPs did indeed form complexes that were detected in the SBA assay. In particular, both GLP1R and GLP2R demonstrated complex formation with all three RAMPs, and GHRHR formed a complex with RAMP2 and RAMP3, but not with RAMP1. These receptors had been judged not to form complexes with RAMPs because over-expression of the receptors in HEK293 or COS-7 cells did not cause co-expressed RAMPs to translocate to the cell surface *(9)*. Our results show directly the existence of GPCR-RAMP complexes and suggests that RAMP translocation studies may not be sensitive to detect all GPCR-RAMP interactions *(23)*. The method we present detects interactions occurring anywhere in the cells, as opposed to just the cell membrane. Of note, preliminary data from another report did suggest that RAMPs were detected at the cell surface when co-expressed with GLP2R in HEK293T cells *(24)*.

Interactions between RAMPs and GPCRs beyond the secretin-like family have remained largely unexplored, with the exception of GPER30 and CaSR *(5, 6, 25)*. Here, among the few non-secretin-like receptors targeted by our SBA, we discovered interactions of RAMPs with chemokine receptors and orphan receptors. The orphan receptors GPR4 and GPR182 interacted with all three RAMPs. Intriguingly, GPR4 demonstrates cell-type selectivity in response to lysolipids, potentially a result of differential RAMP expression in different cell types *(26)*. The chemokine receptors ACKR3/CXCR7 and CXCR3 also interacted weakly with RAMP2 and RAMP3.

Notably, two of the GPCRs that were demonstrated to interact with RAMP2 and RAMP3 have been linked to ligands targeting CALCRL in complex with one of the three RAMPs. These GPCRS, ACKR3/CXCR7 and GPR182, have not been previously reported to interact with RAMPs. GPR182 has been reported to be a receptor for adrenomedullin, a peptide ligand that signals through the CALCRL/RAMP2 complex *(27)*. GPR182 was reclassified as an orphan receptor when these results could not be reproduced *(28)*. ACKR3/CXCR7, a chemokine receptor, was originally described as a receptor for both adrenomedullin and calcitonin-gene related (CGRP), a peptide ligand that signals through the CALCRL/RAMP1 complex *(29)*. More recently, ACKR3/CXCR7 was demonstrated to act as decoy receptor for adrenomedullin *(30)*. This opens up the possibility that RAMP2 can facilitate adrenomedullin or CGRP binding to a receptor.

Recent structural studies provide insight into the potential role of GPCR-RAMP complex formation. A cryo-EM structure of the CALCRL-RAMP1 complex shows that RAMP1 forms extensive contacts with transmembrane helices 3, 4 and 5 as well as with extracellular loop 2 (ECL2) of CALCRL *(31)*. The agonist ligand of CALCRL-RAMP1 makes contacts with the with ECL2 of CALCRL. In contrast, there are very minimal direct contacts between the ligand and RAMP1, suggesting that the complex should exist in the absence of ligand *(32)*. The extensive contact surface between CALCRL and RAMP1 suggests that the complex should be stable under conditions used here in the SBA assay. Given that the CALCRL-RAMP1 structure is generalizable to other possible GPCR-RAMP complexes, Abs that would capture the GPCR through ECL2 might fail to recognize the same GPCR in complex with a RAMP. In line with this hypothesis, we found that a subset of validated anti-GPCR Abs were unable to capture a GPCR-RAMP complex that were identified through capture using other Abs.

We validated a set of Abs to detect GPCR-RAMP complexes in order to avoid artifacts inherent in having to rely upon insufficiently characterized Abs *(33, 34)*. Cross-reactivity of anti-GPCR Abs is a particular problem due to structural and amino acid sequence similarity among GPCRs *(35)*. The multiplexed nature of the SBA assay allowed us to both validate Abs to the RAMPs and GPCRs and check for potential cross-reactivity and off-target binding to all other GPCRs included in the study. We assessed Ab cross-reactivity even in the absence of the intended GPCR while unintended GPCR targets were over-expressed. Using the SBA, we found at least one Ab that targeted the intended GPCR with high selectivity for 19 of the 21 targets with HPA Abs available. We also found at least one Ab to each RAMP that specifically captured the targeted RAMP. Future experiments will validate additional anti-GPCR Abs as they become available. In addition, the SBA strategy can be used to screen for therapeutic mAbs that show the lowest cross-reactivity with other GPCRs.

By being able to use protein-specific, anti-GPCR Abs in the multiplexed SBA assay, the requirement to use engineered epitope tags and a heterologous overexpression system would be eliminated. This would alleviate the potential of identifying non-physiological interactions, since the requirement for overexpressing both proteins would be removed. Thus, we determined whether anti-GPCR Abs can capture the GPCR-RAMP complexes directly. The pre-validated anti-GPCR Abs used in the SBA were able to capture the majority of the GPCR-RAMP complexes as demonstrated by using OLLAS mAb detection.

To our knowledge, no robust proteomics approaches are available to address GPCR-protein interactions. GPCRs are underrepresented in mass spectrometry data, in part due to the lack of efficient proteolysis procedures *(36)*. Affinity assays to screen for GPCR-protein interactions, or membrane protein-protein interactions in general, have been lacking due to difficulties in (i) generating Abs that are functional in the intended assay system *(35, 37)* and (ii) extracting GPCRs from the native environment while maintaining both their binding capabilities and their accessibility for Ab-based detection. Given that at least one Ab to 19 of the 21 GPCRs with Abs in the HPA library was validated for the SBA assay system, it is feasible that a far more complete set GPCRs can be captured by an SBA-based assay once additional GPCR Abs and reagent sources are investigated. It is important to note that many additional anti-GPCR Abs, validated by other immunoassays, became available from HPA following the performance of the experiments described here.

In order to show the presence of selected GPCR-RAMP complexes in membranes, we used PLA, which is an immunolocalization assay in which the proximity of two different Abs is detected using oligonucleotide-labeled secondary Abs. The distance constraints for PLA proximity detection are fairly stringent and the fluorescence signal is highly amplified using a rolling circle DNA polymerization that hybridizes with fluorescent complementary oligonucleotides. The PLA can be carried out *in situ* in a cell membrane environment, which provides evidence for the functional relevance of the SBA results. While PLA has been used to detect GPCR heteromers *(38)*, to our knowledge there have been no reports on its use to detect GPCR-RAMP complexes. Cells co-transfected with CALCRL+RAMP2 and treated with both primary Abs showed a significant (P<0.0001) PLA signal against all the negative controls, indicating that the assay is robust. We also compared results from the PLA using methanol-fixed cells to those using FA-fixed cells. The difference between the methanol and FA data sets was not significant, ensuring that most of the signal came from protein complexes on the membrane.

Although transient protein overexpression by co-transfection is subject to experimental variation, PLA puncta can be quantified, which together allows a semi-quantitative assessment of the extent of GPCR-RAMP interactions within the cellular context. There was a range of GPCR-RAMP interactions as reflected in the number of puncta detected for each GPCR-RAMP2 pair subjected to co-transfection and PLA. For example, co-expression of CXCR3 and RAMP2 did not result in a significant number of puncta compared with mock-transfected control cells. The weak interaction can explain why CXCR3-RAMP2 was not detected through N-termini epitope tags by the lysate-based SBA. The PTH1R-, GPR182-, and GPR4-RAMP2 complexes resulted in a number of puncta that was intermediate between the negative control and the CALCRL-RAMP2 complex.

In summary, the multiplexed SBA assay we developed enables the validation and use of Abs for the detection of GPCR-RAMP interactions. As a proof-of-concept, we defined the complete interactome between secretin-family GPCRs and RAMPs (Fig. 4). We also identified a number of novel GPCR-RAMP interactions, including some interacting partners among rhodopsin-family GPCRs that we selected for study based on earlier bioinformatics analysis. We used the *in situ* PLA to verify the extent of the interaction between several GPCR-RAMP2 pairs tested in the SBA. The SBA is scalable, allowing up to 500 beads to be conjugated with different Abs and decoded in the Luminex system. Therefore, we intend to expand the existing SBA, which consisted of 70 beads, based on the validation of additional Abs using epitope-tagged controls. We also look forward to using the SBA strategy to study other protein-protein interactions, including, but not limited to, GPCR heterodimerization. We consider the SBA technology to be potentially transformative with respect to studies of membrane protein systems.

## Methods and Materials

Study Design: The objective of this study was to identify GPCR-RAMP complexes and validate Abs against each GPCR and each RAMP. Research subjects were detergent solubilized lysates from transfected HEK293 Freestyle cells. For each GPCR-RAMP combination, lysates were prepared from at least three different co-transfections performed on different days. Each prepared lysate was analyzed by the SBA in duplicate. The treatments applied were different GPCR and RAMPs transfected in a controlled laboratory design. Median fluorescence intensity (MFI) of at least 20 events per bead ID was used for data processing. For the PLA experiments, two biological replicates were performed for the negative control experiments that included only one primary Ab, and at least three biological replicates were performed for each GPCR-RAMP pair.

### Materials

1D4 and OLLAS Abs were conjugated to PE using an Ab conjugation kit from Abcam according to manufacturer’s protocols. Information on the source of all Abs used in the SBA can be found in table S1. HPA Abs from the Human Protein Atlas are commercially available from Atlas Antibodies AB. HEK293 Freestyle cells (HEK293F), Freestyle 293 Expression media, Freestyle Max Reagent, and 125 mL culture flasks were from Thermo Fisher. Phycoerythrin (PE) conjugated HA and FLAG Abs, as well as unconjugated HA Ab, were from Biolegend. 1D4 and OLLAS Abs were conjugated to PE using an Ab conjugation kit from Abcam (Cambridge, MA) according to manufacturer’s protocols. Half volume, 96-well plates were from Grenier (Tucson, AZ). Blocking reagent for ELISA (BRE) and cOmplete protease inhibitor tablets were from Roche (Basel, Switzerland). Phosphate buffered saline (PBS) was from Medicago. ProClin 300, casein, polyvinyl alcohol (PVA), and polyvinylpyrrolidone (PVP), and FLAG M2 Ab were from Sigma (St. Louis, MO). Dodecyl-D-maltopyranoside (DM) was from Anatrace. Rabbit-IgG was from Bethyl. Expression constructs of human GPR182 and GPR4 were obtained from cDNA.org. Anti-mouse IgG and anti-rabbit IgG conjugated R-phycoerythrin (PE) were from Jackson Immuno Research.

Cell culture and transfection: Transient transfections were performed using Freestyle Max Reagent (Thermo Fisher) according to manufacturer’s instructions. HEK293F cells were cultured in serum-free Freestyle 293 Expression media using 125 mL disposable culture flasks (Thermo Fisher). Cells were shaken constantly at 125 rpm at 37°C with 5% CO2. The day prior to transfection cells were diluted to 600,000 cells/mL and allowed to grow overnight. The next day 3 mL of cells were transferred to one well of a 6-well plate. Each well of cells was transfected with 0.5 µg indicated RAMP plasmid DNA, and/or 0.5 µg indicated GPCR plasmid DNA with 3 µL Freestyle MAX Reagent. Total transfected plasmid DNA was kept constant at 3 µg by adding empty vector pcDNA3.1(+).

### DNA constructs

Epitope-tagged human GPCR and RAMP DNA constructs were encoded in a pcDNA3.1(+) mammalian expression vector. The human RAMP1, RAMP2 and RAMP3 cDNAs encoded an N-terminal FLAG tag (DYKDDDDK) following the signal peptide (amino acids 1-27, 1-42 and 1-27 for RAMP1, RAMP2 and RAMP3 respectively) and a C-terminal OLLAS tag (SGFANELGPRLMGK). The human secretin-like GPCR and ADGRF5 cDNAs encoded an 5-hydroxytryptamine receptor 3a receptor (5-HT3a) signal sequence (MRLCIPQVLLALFLSMLTGPGEG) in place of the native signal sequence, as determined by SignalP 4.1 (see Table below) *(39)*. In addition, an N-terminal HA tag (YPYDVPDYA) followed the 5HT3a signal sequence and the C-terminus encoded a 1D4 tag (TETSQVAPA). The RAMP and secretin-like GPCR DNA constructs were codon optimized for expression in human cell lines. Expression constructs of human GPR182 and GPR4 cDNAs encoded an N-terminal HA tag (cDNA.org). The human CCR5, CCR7 and CXCR4 cDNA encoded a C-terminal 1D4 epitope tag. Epitope tags on each of the GPCR DNA constructs are summarized in the table below.

| GPCR | Signal Peptide |
| --- | --- |
| ADCYAP1R1 | 1-23 |
| ADGRF5 | 1-21 |
| CALCRL | 1-22 |
| CALCR | 1-42 |
| CRHR1 | 1-23 |
| CRHR2 | 1-19 |
| GIPR | 1-25 |
| GCGR | 1-26 |
| GLP1R | 1-23 |
| GLP2R | None |
| GHRHR | 1-22 |
| PTH1R | 1-28 |
| PTH2R | 1-24 |
| SCTR | 1-27 |
| VIPR1 | 1-30 |
| VIPR2 | 1-20 |

### Lysate Preparation

Cells were solubilized with dodecyl maltoside (DM) detergent (Anatrace) to form micelles around membrane proteins and maintain GPCR and RAMP structure and complex formation. Twenty-four hours post-transfection, transfected HEK293F cells were harvested and washed once with cold phosphate buffered saline (PBS). Cells were then incubated in solubilization buffer (50 mM HEPES, 1 mM EDTA, 150 mM NaCl, 5 mM MgCl2, pH 7.4) with 1% (w/v) DM and complete protease inhibitor (Roche) for two hours at 4°C with nutation. Following solubilization, lysates were clarified by centrifugation at 22k x g for 20 min at 4°C. Lysates were then transferred to a microcentrifuge tube and total protein content was determined by Protein DC assay according to manufacturer’s specifications (Bio Rad).

### Generation of Suspension Bead Array

The SBA was produced by covalently coupling Abs to MagPlex Beads (Luminex), which are magnetic, bar-coded beads where each bar-code is defined by a unique combination of infrared and near red dyes. Each Ab was coupled to a unique bead identity as previously described *(16)*. In brief, 1.6 µg of each Ab was diluted in MES buffer (100 mM 2-(N-morpholino)ethanesulfonic acid (MES) pH 5.0) to a final volume of 100 µL. The carboxylated surface of the magnetic beads was then activated with N-hydroxysuccinimide (Pierce) and 1-Ethyl-3-(3-dimethylaminopropyl)-carbodiimide (Proteochem). After twenty minutes incubation with the activation solution, the diluted Abs were added to the beads and coupling occurred for two hours at room temperature. Beads were then washed and stored in a protein-containing blocking buffer, BRE (Roche), with the addition of ProClin300 (Sigma). Coupling efficiency for each Ab was determined by incubating the beads with PE-conjugated anti-rabbit IgG or PE-conjugated anti-mouse IgG (Jackson Immuno).

SBA assay procedure: Lysates were incubated with an aliquot of the SBA and protein association with each bead was detected with a PE-conjugated Ab. Lysates were diluted to 50 µg/mL in solubilization buffer containing 0.1% (w/v) DM, to a total volume of 25 µL in a half-volume, 96-well plate. An equal amount of 2x SBA assay buffer (PBS containing 1% (w/v) PVA, 1.6% (w/v) PVP, 0.2% (w/v) casein, and 20% rabbit-IgG) was added to the diluted lysates. Aliquots of the SBA, containing 50-100 beads for each bead ID, were added to the lysates in 50 µL of 1x SBA assay buffer (PBS containing 0.5% PVA (w/v), 0.8% PVP (w/v), 0.1% (w/v) casein, and 10% rabbit IgG). The lysates and beads were incubated overnight at 4°C. Then the beads were washed five times with 100 µL PBS containing 0.05% Tween-20 (PBST) using a BioTek EL406 washer. Beads were then incubated with 50 µL of PE-conjugated detection Abs diluted in BRE containing 0.1% DM, 0.1% Tween-20 and 10% Rabbit IgG. The final dilutions used for the detection Abs were 1:1000 for PE-conjugated anti-FLAG (Biolegend) and PE-conjugated anti-1D4, 1:500 for PE-conjugated anti-OLLAS, and 1:200 for PE-conjugated anti-HA (Biolegend). Following incubation at 4°C for one hour, the beads were wash three times with 100 µL PBST. After the final wash, 100 µL of PBST was added to the beads and the fluorescence associated with each bead was measured in an FlexMap3D instrument (Luminex Corp).

Proximity ligation assay (PLA): PLA was used to determine the presence of GPCR-RAMP complexes in HEK293T cells co-transfected with a subset of GPCRs and RAMP2 that were chosen based on SBA results. Gelatin-coated coverslips were placed within the wells of a 6-well dish with one well per transfection condition. HEK293T cells were seeded at 100,000 cells/mL and allowed to grow for 24 hours before transfection. The cells were transfected with 0.4 μg (except in one experiment where 0.5 μg was used) of pcDNA3.1(+) vector encoding the GPCR and RAMP2 constructs (see DNA constructs in Methods and Materials for more information). The total amount of DNA used for each transfection was brought to 2 μg with empty pcDNA3.1(+) vector. For the mock transfection, 2 μg of empty pcDNA3.1(+) vector was used. Transfection was performed with 4 μL Lipofectamine 2000 per well of the 6-well dish. The total volume of media per well was maintained at 2 mL. After 24 hours, cells were washed twice in PBS, fixed and permeabilized with ice cold methanol for 5 mins at −20°C and then washed three times with PBS. For the FA (formaldehyde) fixation, cells were fixed for 10 mins at RT with 4% weight:volume FA (Polysciences, Inc) in 1x PBS, final concentration. After fixation, cells were washed three times in PBS and then processed following manufacturer’s instructions for DuoLink In Situ Detection Reagents Red Mouse/Rabbit (DUO92101) using rabbit anti-HA (Cell Signaling Technology) and mouse anti-FLAG (Sigma-Aldrich) primary Abs. After PLA processing, cells were mounted in DuoLink mounting medium with DAPI (Sigma-Aldrich), allowed to incubate at RT, stored overnight at −20°C, and imaged the following day.

### PLA Image Acquisition

Deconvoluted PLA images were acquired with a DeltaVision Image Restoration Inverted Olympus IX-71 Microscope using a 100x oil immersion objective. Excitation/emission wavelengths were 390/435nm for the blue channel (DAPI) and 575/632nm for the red channel (PLA puncta). Exposure times and transmittance percentages were held constant while imaging all samples within the same experiment. At least three Z-stack images (0.2 μm thickness per slice) of different fields of view were captured per coverslip for each control, and at least five Z-stack images were captured for all other samples. Images from maximum projection of Z-stack images and Imaris spot analysis result snapshots are shown in main text figure. The final figure was made in Adobe Illustrator CC.

### Data analysis of PLA

Image processing was done in ImageJ (adding scale bars, generating maximum projections) and Imaris. Nuclei stained with DAPI were counted to obtain total number of cells per image. The PLA puncta were counted in a 3D rendering of each Z-stack in Imaris using the Spot tool. The same Spot parameters (estimated puncta XY and Z diameter, threshold) were used for all samples in all experiments. The puncta count value for each Z-stack was divided by the total number of cells per image and results were plotted in Prism 8 (Graphpad).

### Statistical Analysis

Statistical parameters, including the number of sampled units, N, and methods used for conducting statistical tests are indicated in the figure legends, table legends and described in the results. For Fig. 2, MFI from at least three experiments performed in duplicate were used in statistical analysis. An ordinary one-way ANOVA, with Dunnett’s multiple comparisons test was used to calculate statistical significance (table S2) (Prism 7, Graphpad). Statistical significance of RAMP Ab validation (fig. S3) was determined using a Kruskal-Wallis ANOVA, with at least 200 experiments performed in duplicate (R package). Statistical significance of GPCR Ab validation, and GPCR-RAMP complex detection captured by anti-GPCR Abs (Fig. 3 and figs. S4 and S5) was determined using a single-tail Z-test comparing signal from all Abs for each lysate, with at least three experiments performed in duplicate (Excel, Microsoft). A one-way ANOVA followed by Dunnett’s multiple comparisons test was used to compare the mean PLA puncta count of the positive condition (CALCRL+RAMP2, both primary Abs) to that of each of the controls, and to compare CXCR3-RAMP2 and Mock to the other GPCR-RAMP2 pairs (Fig. 5) (Prism 8, Graphpad). Outliers from each GPCR-RAMP pair were determined in Prism via the ROUT method with Q=1%. Two outliers were removed from the PTH1R-RAMP2 data set and three from the CXCR3-RAMP2 data set (Fig. 5). A two-tailed P-test in Prism was used to compare the numbers of PLA puncta from methanol-fixed and PFA-fixed cells (fig. S7). Significance was determined by P<0.05. The alpha level used for each test was 0.05.

## Supplementary Materials

### Supplementary Methods

Fig. S1. Co-expression of GPCR clusters with RAMPs and the position of selected GPCRs on the phylogenetic tree.

Fig. S2. Validation of epitope tag Abs to capture and detect RAMPs and GPCRs.

Fig. S3. Validation of Abs used to capture RAMPs.

Fig. S4. Analysis of anti-GPCR Ab cross-reactivity.

Fig. S5. Detection of GPCR-RAMP complexes following capture by all anti-GPCR Abs.

Fig. S6. Statistical validation of GPCR-RAMP SBA data sets.

Fig. S7. Detection of CALCRL-RAMP2 interactions in cell membranes using PLA.

Table S1. The ID of the bead coupled to each specific Ab, the source of the Ab, and product code.

Table S2. Statistical significance of GPCR-RAMP complex formation using epitope tags for capture.

Table S3. Overall statistic for GPCR-RAMP complex formation.

## Acknowledgments

The authors would like to thank Maja Neiman for experimental input, the Bio-Imaging Resource Center (BIRC) at Rockefeller University for advice and access to instrumentation, Mun-Gwan Hong, Cecilia Engel Thomas and Don Lewis for discussion concerning data analysis. The KTH Center for Applied Precision Medicine (KCAP), funded by the Erling-Persson Family Foundation, is acknowledged for financial support. This work was supported by grants for Science for Life Laboratory, as well as the Knut and Alice Wallenberg Foundation for funding the Human Protein Atlas. Travel for this research was supported by the Nicholson Short-Term Exchange, Albert Cass Fellowship, and Alexander Mauro Fellowship.

## Author Contributions

E.L., I.K., T.H., J.S., and T.S. designed experiments. E.L., I.K., E.P. and E.C. performed experiments. T.D.C., E.L., I.K., R.V., and J.S. analyzed the data. J.S., T.H. and T.S. supervised the project. E.L., I.K., T.H., J.S., and T.S. wrote the manuscript.

## Data and materials availability

Data and materials are available upon request.

