## Supplementary Materials for "Multiplexed Analysis of the Secretin-like GPCR-RAMP Interactome"

**Supplementary Methods**

Co-expression analysis: RNA-seq data from 53 tissues provided by 544 donors, with a total of 8555 samples, were downloaded from gtexportal.org (GTEx_Analysis_v6p_RNA-seq_RNA-SeQCv1.1.8_gene_rpkm.gct.gz) *(41)*. Downloaded data was provided as reads per kilobase per million mapped reads (RPKM). Only samples passing quality control were included in the dataset. Read counts and RPKM values were produced with RNA-SeqC; importantly, reads were mapped to a single gene (see gtexportal.org documentation for more information). For quantile normalization, gene expression across an individual sample was fit to the averaged distribution observed across samples. Prior to implementing spearman correlation analysis, the median normalized expression per tissue was calculated to account for differences in the number of samples per tissue type. Spearman correlation coefficient was calculated for each RAMP/non-olfactory GPCR pair. GPCR clusters were assigned as per Fredriksson et al *(42)*.

Statistical comparison of data sets: The data obtained using anti-GPCR capture Abs was compared to the data obtained using the epitope-tag capture methods. Based upon the data set for GPCR-RAMP complexes derived from the epitope-tag capture Abs, a matrix of hypothetical outcomes for the data set derived from the anti-GPCR Ab capture strategy was constructed. To compare the two matrices, the results were converted to binary score matrices (0,1) and the two matrices (epitope-tag Ab capture *versus* direct Ab capture). A Z-score threshold of 1.645 was applied for the anti-GPCR Ab data set of 1.645, which corresponds to a confidence interval of 95% for a single-tailed test (fig S5). The threshold used to convert the summarized and normalized epitope tag data to binary form was increased by an interval of 0.125 (table S3). The following metrics were plotted as a function of the threshold used to convert the epitope tag data to a binary matrix: (1) overall percent agreement ($P_0$), (2) the percent of hits from the epitope tag data that are also found in the anti-GPCR Ab data (sensitivity), (3) the percent of non-hits from the epitope tag data that are also non-hits in the anti-GPCR Ab data (specificity), (4) the probability of a positive result in the anti-GPCR Ab data also being a positive in the epitope tag data (positive predictive value), (5) the probability of a negative result in the anti-GPCR Ab data also being a negative in the epitope tag data (negative predictive value) and (6) the similarity of the two data sets (Jaccard Index). The formulas used are listed below (TP, true positive; TN, true negative; FP, false positive; TP, true positive). See the table below for formulas used.

| Metric | Formula | Information |
|---|---|---|
| $P_o$ | (TP+TN)/(TP+TN+FP+FN) | Overall percent agreement |
| Sensitivity | TP/(TP+FN) | Percent of all positives from epitope beads that the anti-GPCR beads detect |
| Specificity | TN/(TN+FP) | Percent of all negatives from epitope beads that the anti-GPCR beads detect |
| Positive Predictive Value | TP/(TP+FP) | Probability of being a positive from epitope beads if anti-GPCR beads says positive |
| Negative Predictive Value | TN/(TN+FN) | Probability of being a negative from epitope beads if anti-GPCR beads says negative |
| Jaccard Index | TP/(TP+FP+FN) | Similarity of positives between the epitope beads and the anti-GPCR beads |

### Supplementary Figures

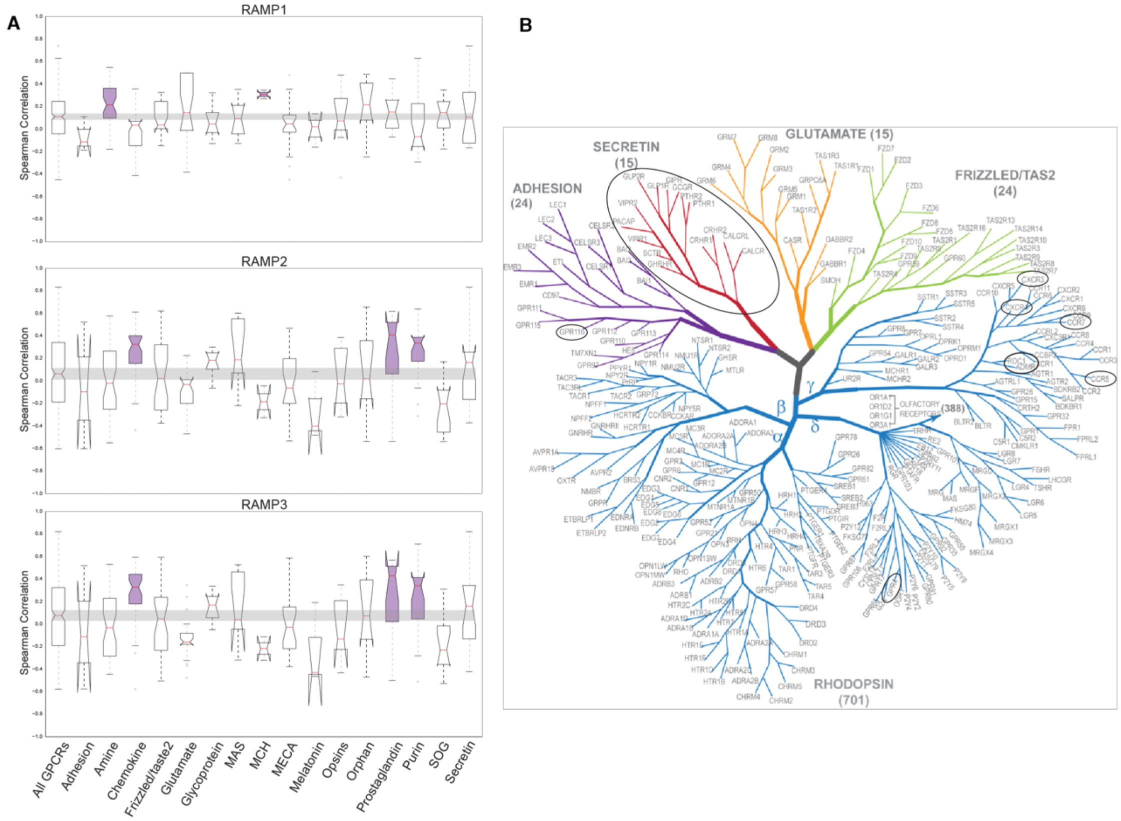


**Fig S1.** Co-expression of GPCR clusters with RAMPs and the position of selected GPCRs on the phylogenetic tree. **(A)** Co-expression of GPCRs clusters with each RAMP in comparison to all GPCRs. Boxplot of spearman correlation coefficients across 53 human tissues between RAMP-GPCR pairs. Notches indicate 95% confidence intervals of the median and the grey bars indicate the 95% confidence interval of the median for all GPCRs. Receptor clusters with significantly higher co-expression than all GPCRs are highlighted with purple. Abbreviations: MCH, melanin-concentrating hormone; MECA, melanocortin, endothelial differentiation, cannabinoid, and adenosine; SOG, somatostatin, opioid, and galanin. **(B)** GPCR phylogenetic tree highlighting the receptors tested for complex formation with RAMPs (Figure adapted from *(43)*). Circled GPCR names indicate that the receptor was included in this study. In the phylogenetic tree, ACKR3 is referred to as RDC1, GPR182 is ADMR, ADGRF5 is GPR116, and ADCYAP1R1 is PACAP.

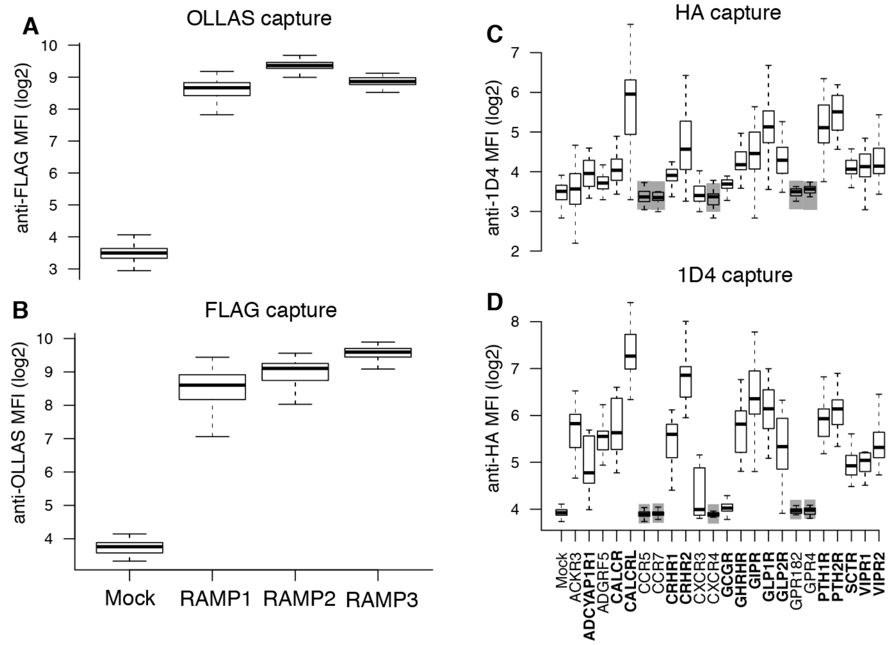


**Fig. S2.** Validation of epitope tag Abs to capture and detect RAMPs and GPCRs. Lysates from cells transfected with each epitope-tagged RAMP construct (FLAG and OLLAS) were incubated with the SBA, which included mAbs targeting FLAG or OLLAS. Each RAMP was **(A)** captured with anti-OLLAS mAb bead and detected with PE-conjugated anti-FLAG mAb or **(B)** captured with anti-FLAG mAb bead and detected with a PE-conjugated anti-OLLAS mAb. Lysates from cells transfected with each epitope-tagged GPCR construct (HA and/or 1D4) were incubated with the SBA, which included mAbs targeting HA or 1D4. Each GPCR was **(C)** captured with anti-HA mAb bead and detected with a PE-conjugated anti-1D4 mAb, or **(D)** captured with anti-1D4 mAb bead and detected with a PE-conjugated anti-HA mAb. Grey boxes around the occasional data set indicates that the GPCR does not have both engineered epitope tags, and thus would not be expected to show signal in this experiment. The labels in bold correspond to secretin-like GPCRs. Data is median fluorescence intensity (MFI) and representative of at least nine experiments performed in duplicate. Validated Abs are underlined.

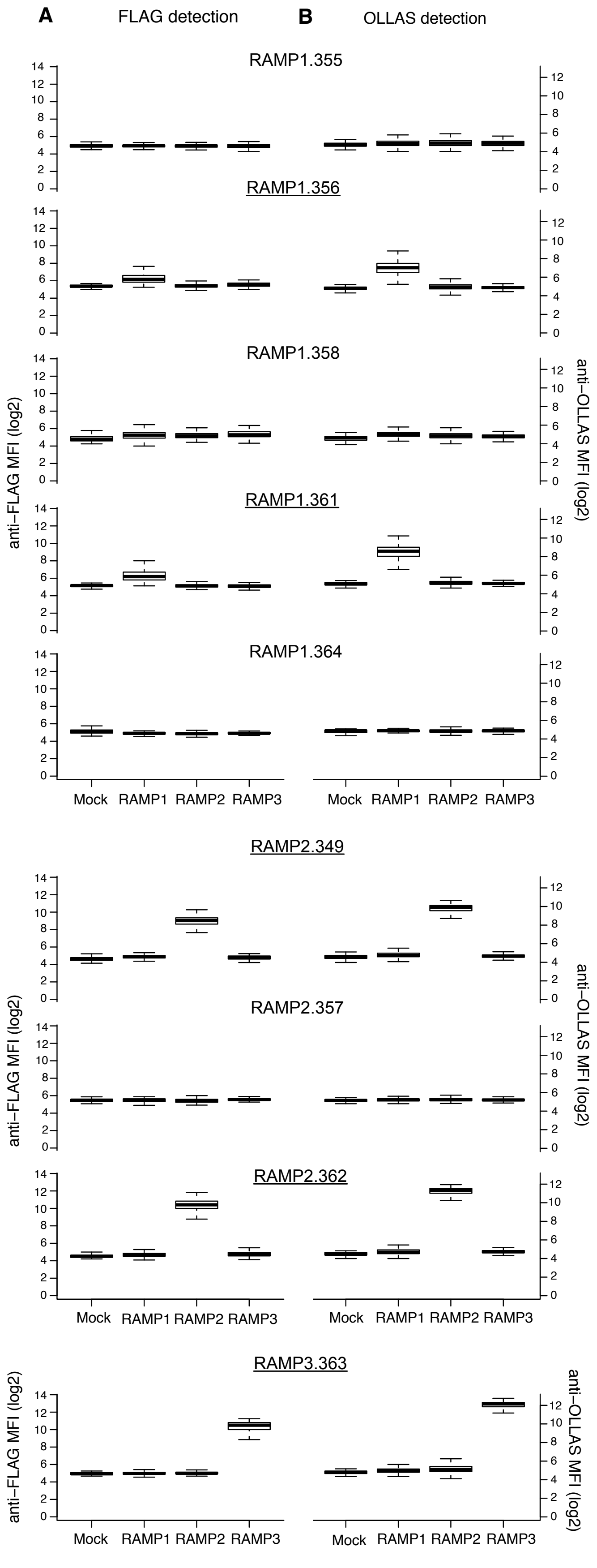
A
FLAG detection
B
OLLAS detection
RAMP1.355
RAMP1.356
RAMP1.358
RAMP1.361
RAMP1.364
anti-FLAG MFI (log2)
anti-OLLAS MFI (log2)
Mock
RAMP1
RAMP2
RAMP3
RAMP2.349
RAMP2.357
RAMP2.362
RAMP3.363

64

**Fig. S3.** Validation of Abs used to capture RAMPs. In order to validate anti-RAMP Abs, lysates from cells transfected with each epitope-tagged RAMP construct (FLAG and OLLAS) were incubated with the SBA, which included beads conjugated with nine capture Abs targeting the three RAMPs. **(A)** PE-conjugated anti-FLAG and **(B)** PE-conjugated anti-OLLAS mAbs were used to detect any RAMPs captured by the beads. Data are median fluorescence intensity (MFI) and represent at least 200 experiments, each performed in duplicate. At a statistical significance of $p \leq 0.05$ (Kruskal-Wallis ANOVA), we validated at least one capture Ab for each of the three RAMP s (a total of 5 RAMP capture Abs). Bead ID numbers are listed after each RAMP name and the corresponding Ab name is provided in table S1.

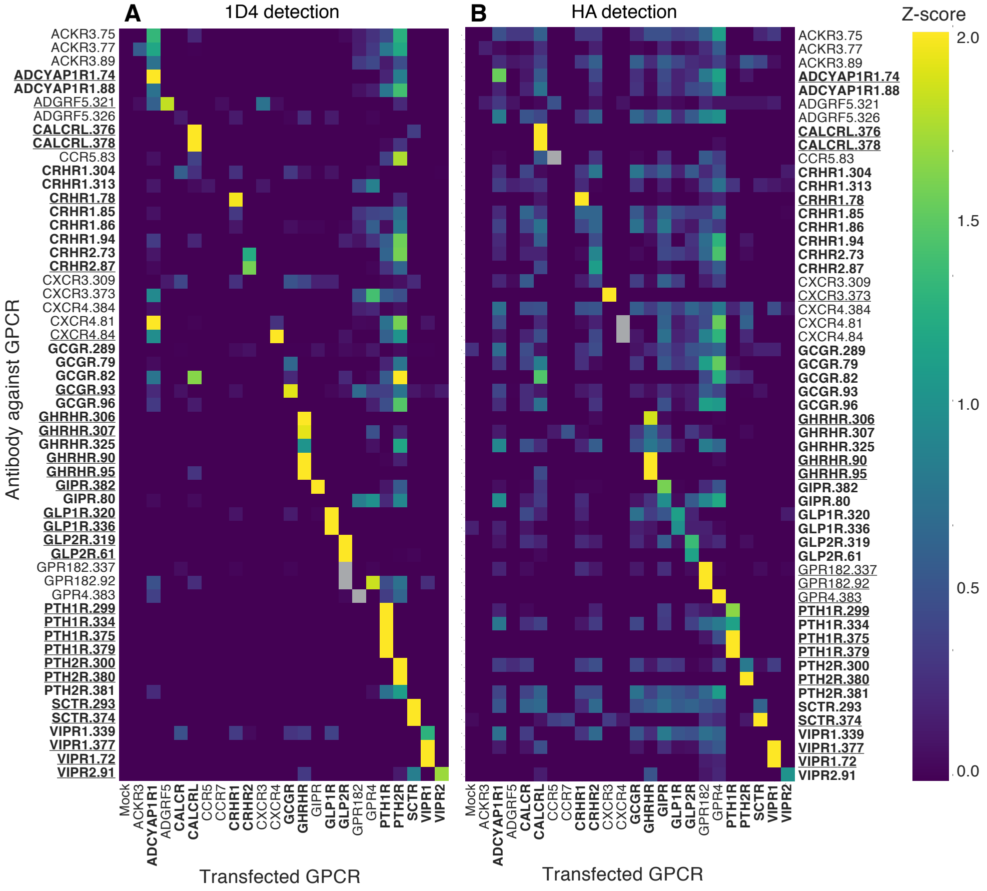


**Fig. S4.** Analysis of anti-GPCR Ab cross-reactivity. Lysates from cells transfected with each epitope-tagged GPCR construct (HA and 1D4) were incubated with the SBA, which included 55 Abs to 21 GPCRs. **(A)** PE-conjugated anti-1D4 and **(B)** PE-conjugated anti-HA were used to detect any GPCRs captured by the beads. The occasional grey boxes indicate that the GPCR did not have the appropriate epitope tag to be detected. The labels in bold correspond to secretin-like GPCRs and validated Abs are underlined Abs. Heatmaps represent the z-scores of median fluorescence intensity (MFI) and indicates the ability of the GPCR Abs to capture each of the 23 GPCRs used in the study. Data represents the median z-score of at least three experiments performed in duplicate. Bead ID numbers are listed after each GPCR name and the corresponding Ab name is provided in table S1.

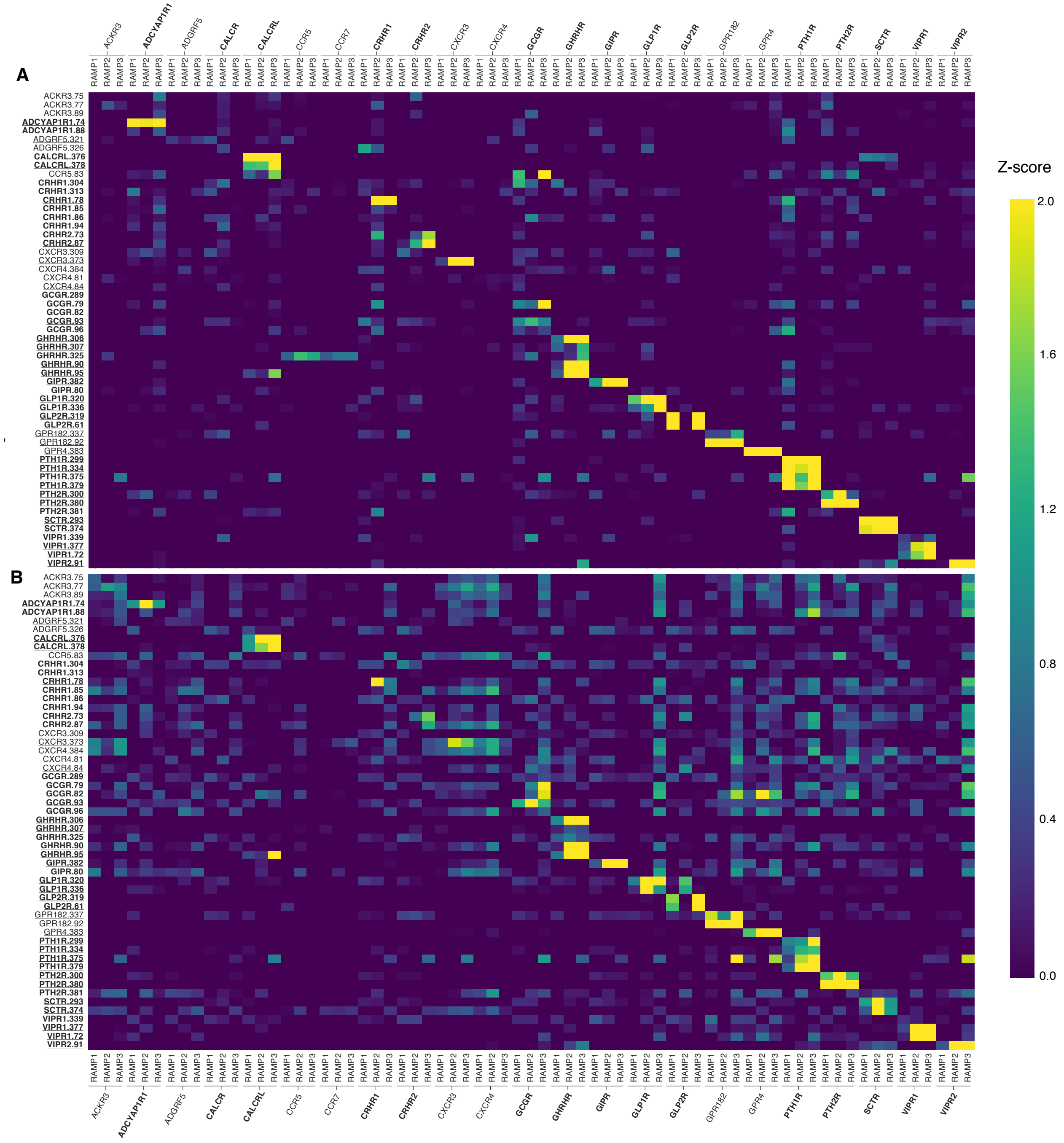


**Fig. S5.** Detection of GPCR-RAMP complexes following capture by all anti-GPCR Abs. Lysates from cells transfected with each epitope-tagged RAMP construct (FLAG and OLLAS) and co-transfected with each epitope-tagged GPCR construct (HA and 1D4) were incubated with the SBA, which included 55 Abs to the 21 GPCRs included in this study. Complexes were captured in multiplex fashion using the anti-GPCR Ab beads. The presence of a GPCR-RAMP complex was detected using **(A)** PE-conjugated OLLAS mAb, or **(B)** PE-conjugated anti-FLAG mAb. The labels in bold correspond to secretin-like GPCRs and validated Abs are underlined. The heatmap displays the Z-score of median fluorescence intensity (MFI) and represents at least three experiments performed in duplicate. Bead ID numbers are listed after each GPCR name and the corresponding Ab name is provided in table S1.

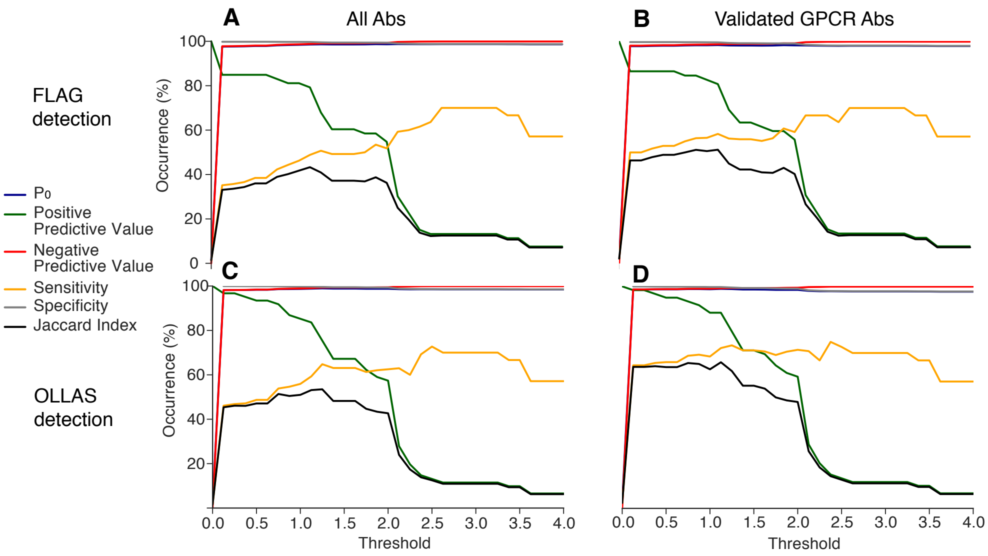


**Fig. S6.** Statistical validation of GPCR-RAMP SBA data sets. Data from the capture of the GPCR-RAMP complexes with anti-epitope mAbs were compared with data obtained from the GPCR-RAMP complexes captured using anti-GPCR Abs. PE-conjugated anti-FLAG was used to detect GPCR-RAMP complexes captured using **(A)** all anti-GPCR Abs or **(B)** validated anti-GPCR Abs. Alternatively, PE-conjugated anti-OLLAS mAb was used to detect GPCR-RAMP complexes captured using **(C)** all anti-GPCR Abs or **(D)** validated anti-GPCR Abs. The Z-score threshold for the anti-GPCR Ab data was set at 1.645. $P_0$ (blue), positive predictive value (green), negative predictive value (red), sensitivity (yellow), specificity (grey), and Jaccard Index (black) are plotted as a function of increasing threshold for the interaction results using epitope tags for capture and detection (Supplemental Table 3). Supplementary Materials and Methods shows the formulas and metrics used and provides a narrative description of each of the statistical terms. For example, the Jaccard Index represents the overall agreement of the positive results in both data sets and indicates at which thresholds the agreement is maximized.

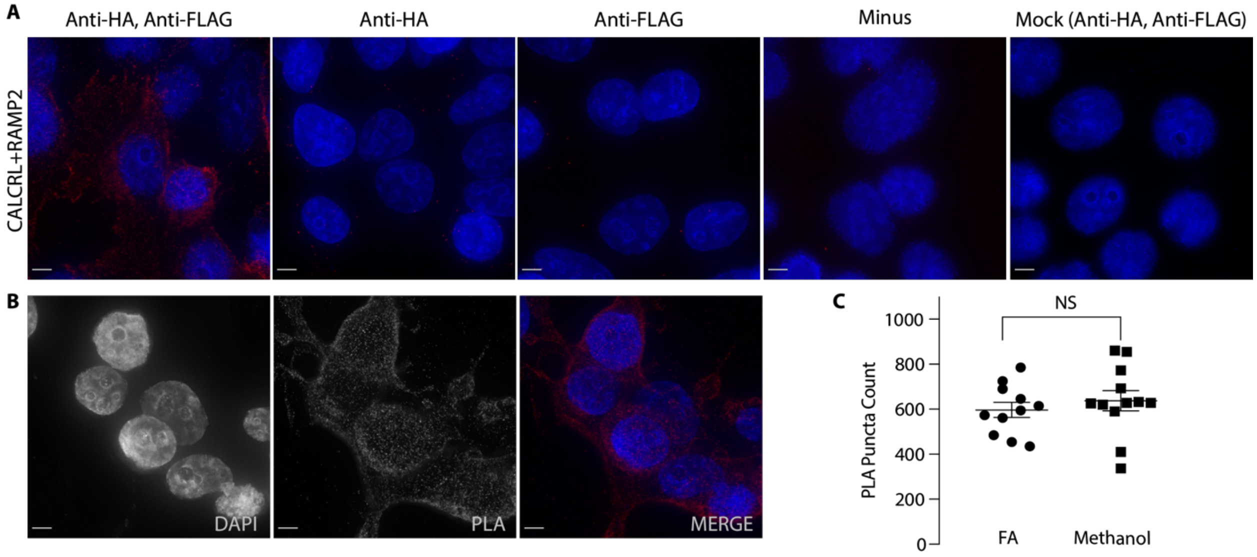


**Fig. S7.** Detection of CALCRL-RAMP2 interactions in cell membranes using PLA. Cells co-transfected with epitope-tagged GPCR and RAMP2 then incubated with anti-HA and anti-FLAG Abs. PLA was then carried out to detect GPCR-RAMP2 interactions. **(A)** Representative images of PLA performed on CALCRL+RAMP2 co-transfected cells using Ab detection as noted. Images show maximum projection of Z-stack, which is the maximum signal intensity for each channel at each point across all slices in the Z-stack. **(B)** Representative PLA images showing greyscale split-channel view of a Z-stack maximum projection for cells co-transfected with CALCRL+RAMP2 and treated with both primary Abs. The merge is presented in color. Scale bars, 5μm for both (A) and (C). Blue = DAPI, red = PLA puncta. **(C)** PLA puncta counts per cell for cells co-transfected with CALCRL+RAMP2, fixed with either FA or methanol and subjected to PLA. Data are from at least two experiments performed with at least five replicates. Significance determined by two-tailed P-test (P=0.4763, NS = not significant).

**Table S1.** The ID of the bead coupled to each specific Ab, the source of the Ab, and product code. The labels in bold correspond to secretin-like GPCRs. Underlined product codes indicate validated Abs.

| Bead ID | Protein Name | Antibody Source | Product Code | Bead ID | Protein Name | Antibody Source | Product Code |
|---|---|---|---|---|---|---|---|
| 61 | **GLP2R** | HPA | HPA027929 | 321 | ADGRF5 | HPA | HPA065251 |
| 72 | **VIPR1** | HPA | HPA046516 | 325 | **GHRHR** | HPA | HPA068576 |
| 73 | **CRHR2** | HPA | HPA046683 | 326 | ADGRF5 | HPA | HPA068796 |
| 74 | **ADCYAP1R1** | HPA | HPA030739 | 334 | **PTH1R** | HPA | HPA075879 |
| 75 | ACKR3 | HPA | HPA049718 | 336 | **GLP1R** | HPA | HPA077988 |
| 77 | ACKR3 | HPA | HPA032003 | 337 | GPR182 | HPA | HPA027037 |
| 78 | **CRHR1** | HPA | HPA063352 | 343 | Empty | N/A | N/A |
| 79 | **GCGR** | HPA | HPA066333 | 339 | **VIPR1** | HPA | HPA026777 |
| 80 | **GIPR** | HPA | HPA068054 | 349 | RAMP2 | HPA | HPA064452 |
| 81 | CXCR4 | HPA | HPA068321 | 350 | rabbit IgG | Bethyl | P120 |
| 82 | **GCGR** | HPA | HPA071228 | 355 | RAMP1 | Abcam | ab156575 |
| 83 | CCR5 | HPA | HPA070587 | 356 | RAMP1 | HPA | HPA010654 |
| 84 | CXCR4 | HPA | HPA051623 | 357 | RAMP2 | HPA | HPA052020 |
| 85 | **CRHR1** | HPA | HPA055287 | 358 | RAMP1 | HPA | HPA057814 |
| 86 | **CRHR1** | HPA | HPA071484 | 359 | OLLAS | In house | N/A |
| 87 | **CRHR2** | HPA | HPA073345 | 360 | FLAG | Sigma | F3165 |
| 88 | **ADCYAP1R1** | HPA | HPA073908 | 361 | RAMP1 | RnD | AF6428 |
| 89 | ACKR3 | HPA | HPA057492 | 362 | RAMP2 | RnD | AF6427 |
| 90 | **GHRHR** | **HPA** | HPA077545 | 363 | RAMP3 | RnD | AF4875 |
| 91 | **VIPR2** | **HPA** | HPA062707 | 364 | RAMP1 | Santa | sc-11379 |
| 92 | **GPR182** | **HPA** | HPA027037 | 365 | mouse | Bio Rad | PMP01X |
| 93 | **GCGR** | **HPA** | HPA057075 | 366 | 1D4 | In house | N/A |
| 94 | **CRHR1** | **HPA** | HPA046066 | 367 | HA | Biolegend | 16B12 |
| 95 | **GHRHR** | **HPA** | HPA070884 | 373 | CXCR3 | HPA | HPA003189 |
| 96 | **GCGR** | **HPA** | HPA074345 | 374 | **SCTR** | HPA | HPA007269 |
| 293 | **SCTR** | **HPA** | HPA007312 | 375 | **PTH1R** | HPA | HPA007491 |
| 299 | **PTH1R** | **HPA** | HPA007978 | 376 | **CALCRL** | HPA | HPA007586 |
| 300 | **PTH2R** | **HPA** | HPA010534 | 377 | **VIPR1** | HPA | HPA007588 |
| 304 | **CRHR1** | **HPA** | HPA032018 | 378 | **CALCRL** | HPA | HPA008070 |
| 306 | **GHRHR** | **HPA** | HPA034645 | 379 | **PTH1R** | HPA | HPA007978 |
| 307 | **GHRHR** | **HPA** | HPA034644 | 380 | **PTH2R** | HPA | HPA010534 |
| 309 | CXCR3 | HPA | HPA045942 | 381 | **PTH2R** | HPA | HPA010655 |
| 313 | **CRHR1** | **HPA** | HPA052441 | 382 | **GIPR** | HPA | HPA017428 |
| 319 | **GLP2R** | **HPA** | HPA064671 | 383 | GPR4 | HPA | HPA019207 |
| 320 | **GLP1R** | **HPA** | HPA065175 | 384 | CXCR4 | HPA | HPA027832 |

**Table S2.** Statistical significance of GPCR-RAMP complex formation using epitope tags for capture. The statistical significance of signal between mock transfected cell lysates and lysates from cells co-transfected with each dual-tagged RAMP construct plus each dual or single-tagged GPCR construct. (Ordinary one-way ANOVA, Dunnett's multiple comparisons test). N/A indicates that the GPCR does not have the epitope tag to be captured or detected with the corresponding Ab. Secretin-like receptors are shown in bold.

| | Interaction with RAMP1 | | | | | | | |
|---|---|---|---|---|---|---|---|---|
| Capture Ab | 1D4 | 1D4 | HA | HA | OLLAS | OLLAS | FLAG | FLAG |
| Detection Ab | FLAG | OLLAS | FLAG | OLLAS | 1D4 | HA | 1D4 | HA |
| ACKR3 | 0.857 | 0.804 | 1.000 | 1.000 | 0.999 | 1.000 | 1.000 | 1.000 |
| **ADCYAP1R1** | 0.204 | 0.025 | 1.000 | 1.000 | 0.001 | 1.000 | 0.000 | 1.000 |
| ADGRF5 | 0.392 | 0.017 | 1.000 | 1.000 | 0.706 | 1.000 | 0.907 | 1.000 |
| **CALCR** | 0.079 | 0.002 | 1.000 | 1.000 | 0.986 | 1.000 | 0.994 | 1.000 |
| **CALCRL** | 0.000 | 0.000 | 0.000 | 0.000 | 0.000 | 0.000 | 0.000 | 0.000 |
| CCR5 | 1.000 | 1.000 | N/A | N/A | 1.000 | N/A | 1.000 | N/A |
| CCR7 | 1.000 | 1.000 | N/A | N/A | 1.000 | N/A | 1.000 | N/A |
| **CRHR1** | 0.999 | 0.999 | 1.000 | 1.000 | 1.000 | 1.000 | 1.000 | 1.000 |
| **CRHR2** | 0.927 | 0.808 | 0.999 | 1.000 | 0.994 | 1.000 | 0.999 | 0.999 |
| CXCR3 | 0.999 | 0.969 | 1.000 | 1.000 | 0.999 | 1.000 | 1.000 | 1.000 |
| CXCR4 | 0.999 | 0.999 | 1.000 | 1.000 | 1.000 | 1.000 | 1.000 | 1.000 |
| **GCGR** | 0.016 | 0.000 | 1.000 | 0.999 | 0.904 | 1.000 | 0.855 | 1.000 |
| **GHRHR** | 0.280 | 0.069 | 0.999 | 1.000 | 0.158 | 1.000 | 0.491 | 0.999 |
| **GIPR** | 0.000 | 0.000 | 0.447 | 0.999 | 0.053 | 0.994 | 0.028 | 0.603 |
| **GLP1R** | 0.000 | 0.000 | 0.170 | 0.821 | 0.000 | 0.999 | 0.000 | 0.884 |
| **GLP2R** | 0.000 | 0.000 | 0.944 | 1.000 | 0.000 | 0.995 | 0.000 | 0.494 |
| GPR182 | N/A | N/A | 0.000 | 0.275 | N/A | 0.023 | N/A | 0.000 |
| GPR4 | N/A | N/A | 0.000 | 0.440 | N/A | 0.003 | N/A | 0.000 |
| **PTH1R** | 0.000 | 0.000 | 0.690 | 0.721 | 0.000 | 0.995 | 0.000 | 0.864 |
| **PTH2R** | 0.000 | 0.000 | 0.005 | 0.302 | 0.000 | 0.879 | 0.000 | 0.244 |
| **SCTR** | 0.032 | 0.000 | 1.000 | 0.999 | 0.000 | 0.975 | 0.000 | 1.000 |
| **VIPR1** | 0.630 | 0.181 | 1.000 | 0.999 | 0.073 | 1.000 | 0.106 | 1.000 |
| **VIPR2** | 0.003 | 0.087 | 0.999 | 1.000 | 0.632 | 1.000 | 0.611 | 1.000 |

| | Interaction with RAMP2 | | | | | | | |
|---|---|---|---|---|---|---|---|---|
| Capture Ab | 1D4 | 1D4 | HA | HA | OLLAS | OLLAS | FLAG | FLAG |
| Detection Ab | FLAG | OLLAS | FLAG | OLLAS | 1D4 | HA | 1D4 | HA |
| ACKR3 | 0.000 | 0.000 | 0.988 | 1.000 | 0.907 | 1.000 | 0.070 | 0.999 |
| **ADCYAP1R1** | 0.060 | 0.021 | 1.000 | 1.000 | 0.062 | 1.000 | 0.000 | 1.000 |
| ADGRF5 | 0.000 | 0.000 | 1.000 | 0.999 | 0.987 | 1.000 | 0.977 | 1.000 |
| **CALCR** | 0.000 | 0.000 | 0.958 | 0.985 | 0.994 | 1.000 | 0.456 | 0.999 |
| **CALCRL** | 0.000 | 0.000 | 0.000 | 0.000 | 0.000 | 0.000 | 0.000 | 0.000 |
| CCR5 | 0.408 | 0.153 | N/A | N/A | 0.999 | N/A | 0.999 | N/A |

| | | | | | | | | |
|---|---|---|---|---|---|---|---|---|
| CCR7 | 0.995 | 0.418 | N/A | N/A | 0.975 | N/A | 0.999 | N/A |
| **CRHR1** | 0.001 | 0.000 | 0.994 | 0.999 | 0.999 | 1.000 | 0.988 | 1.000 |
| **CRHR2** | 0.471 | 0.115 | 0.041 | 1.000 | 0.986 | 1.000 | 0.883 | 0.869 |
| CXCR3 | 0.988 | 0.782 | 0.995 | 0.995 | 1.000 | 1.000 | 1.000 | 0.995 |
| CXCR4 | 0.999 | 0.999 | 1.000 | 1.000 | 1.000 | 1.000 | 1.000 | 1.000 |
| **GCGR** | 0.000 | 0.000 | 1.000 | 1.000 | 0.999 | 1.000 | 0.501 | 1.000 |
| **GHRHR** | 0.000 | 0.000 | 0.709 | 1.000 | 0.956 | 1.000 | 0.210 | 0.999 |
| **GIPR** | 0.000 | 0.000 | 0.000 | 0.021 | 0.000 | 0.941 | 0.000 | 0.816 |
| **GLP1R** | 0.000 | 0.000 | 0.009 | 0.556 | 0.003 | 0.996 | 0.000 | 0.827 |
| **GLP2R** | 0.000 | 0.000 | 0.021 | 0.982 | 0.000 | 0.994 | 0.000 | 0.984 |
| GPR182 | N/A | N/A | 0.000 | 0.000 | N/A | 0.000 | N/A | 0.000 |
| GPR4 | N/A | N/A | 0.000 | 0.000 | N/A | 0.000 | N/A | 0.000 |
| **PTH1R** | 0.000 | 0.000 | 0.095 | 0.966 | 0.000 | 0.989 | 0.000 | 0.712 |
| **PTH2R** | 0.000 | 0.000 | 0.065 | 0.090 | 0.000 | 0.254 | 0.000 | 0.648 |
| **SCTR** | 0.001 | 0.000 | 0.999 | 0.925 | 0.000 | 0.944 | 0.000 | 0.999 |
| **VIPR1** | 0.030 | 0.002 | 0.995 | 0.995 | 0.016 | 1.000 | 0.000 | 0.999 |
| **VIPR2** | 0.000 | 0.000 | 0.995 | 0.984 | 0.167 | 1.000 | 0.002 | 0.999 |

| | Interaction with RAMP3 | | | | | | | |
|---|---|---|---|---|---|---|---|---|
| Capture Ab | 1D4 | 1D4 | HA | HA | OLLAS | OLLAS | FLAG | FLAG |
| Detection Ab | FLAG | OLLAS | FLAG | OLLAS | 1D4 | HA | 1D4 | HA |
| ACKR3 | 0.001 | 0.000 | 1.000 | 1.000 | 0.999 | 0.999 | 0.999 | 0.999 |
| **ADCYAP1R1** | 0.024 | 0.000 | 1.000 | 1.000 | 0.005 | 0.981 | 0.190 | 0.999 |
| ADGRF5 | 0.016 | 0.000 | 1.000 | 1.000 | 0.820 | 0.999 | 0.999 | 1.000 |
| **CALCR** | 0.000 | 0.000 | 1.000 | 0.983 | 0.281 | 0.996 | 0.923 | 0.999 |
| **CALCRL** | 0.000 | 0.000 | 0.000 | 0.063 | 0.000 | 0.000 | 0.000 | 0.000 |
| CCR5 | 0.999 | 0.995 | N/A | N/A | 1.000 | N/A | 1.000 | N/A |
| CCR7 | 0.999 | 0.551 | N/A | N/A | 0.994 | N/A | 0.999 | N/A |
| **CRHR1** | 0.123 | 0.000 | 1.000 | 1.000 | 0.999 | 1.000 | 0.999 | 1.000 |
| **CRHR2** | 0.008 | 0.000 | 0.999 | 0.977 | 0.157 | 0.995 | 0.817 | 0.586 |
| CXCR3 | 0.885 | 0.039 | 0.999 | 0.999 | 1.000 | 1.000 | 1.000 | 1.000 |
| CXCR4 | 1.000 | 0.999 | 1.000 | 1.000 | 1.000 | 1.000 | 1.000 | 1.000 |
| **GCGR** | 0.000 | 0.000 | 1.000 | 1.000 | 0.297 | 0.999 | 0.776 | 1.000 |
| **GHRHR** | 0.000 | 0.000 | 0.999 | 0.999 | 0.017 | 0.925 | 0.147 | 0.740 |
| **GIPR** | 0.000 | 0.000 | 0.961 | 0.999 | 0.016 | 0.063 | 0.193 | 0.736 |
| **GLP1R** | 0.000 | 0.000 | 0.964 | 0.372 | 0.040 | 0.306 | 0.017 | 0.522 |
| **GLP2R** | 0.000 | 0.000 | 1.000 | 0.999 | 0.000 | 0.050 | 0.000 | 0.569 |
| GPR182 | N/A | N/A | 0.000 | 0.000 | N/A | 0.000 | N/A | 0.000 |
| GPR4 | N/A | N/A | 0.000 | 0.000 | N/A | 0.000 | N/A | 0.000 |
| **PTH1R** | 0.000 | 0.000 | 0.809 | 0.633 | 0.000 | 0.451 | 0.000 | 0.034 |
| **PTH2R** | 0.000 | 0.000 | 0.350 | 0.141 | 0.000 | 0.078 | 0.000 | 0.145 |
| **SCTR** | 0.003 | 0.000 | 1.000 | 0.995 | 0.000 | 0.955 | 0.000 | 0.984 |
| **VIPR1** | 0.000 | 0.000 | 0.999 | 0.860 | 0.001 | 0.965 | 0.043 | 0.999 |
| **VIPR2** | 0.000 | 0.000 | 0.933 | 0.926 | 0.000 | 0.862 | 0.001 | 0.911 |

**Table S3.** Overall statistic for GPCR-RAMP complex formation. P-values from table S3 of <0.0001 were assigned 4, <0.001 a 3, <0.01 a 2, and <0.05 a 1. The values were summed and divided by the number of capture and detection pairs that we expected to be capable of measuring the relevant complex. For dual-tagged GPCRs, we divided by eight and for single-tagged GPCRs we divided by four to obtain a normalized value. Secretin-like receptors are shown in bold.

| | RAMP1 | RAMP2 | RAMP3 |
|---|---|---|---|
| ACKR3 | 0 | 1.125 | 0.75 |
| **ADCYAP1R1** | 1 | 0.75 | 0.75 |
| ADGRF5 | 0.125 | 0.875 | 0.625 |
| **CALCR** | 0.25 | 1 | 1 |
| **CALCRL** | 4 | 4 | 3.5 |
| CCR5 | 0 | 0 | 0 |
| CCR7 | 0 | 0 | 0 |
| **CRHR1** | 0 | 0.625 | 0.375 |
| **CRHR2** | 0 | 0 | 0.75 |
| CXCR3 | 0 | 0 | 0.25 |
| CXCR4 | 0 | 0 | 0 |
| **GCGR** | 0.625 | 1 | 2 |
| **GHRHR** | 0 | 2 | 1.125 |
| **GIPR** | 1.125 | 2.375 | 1.125 |
| **GLP1R** | 1.75 | 1.875 | 1.25 |
| **GLP2R** | 2 | 2 | 2 |
| GPR182 | 2.25 | 2.25 | 4 |
| GPR4 | 2.5 | 3.25 | 4 |
| **PTH1R** | 2 | 2 | 2.125 |
| **PTH2R** | 2.25 | 2 | 2.125 |
| **SCTR** | 1.625 | 2 | 1.75 |
| **VIPR1** | 0 | 1.125 | 1.25 |
| **VIPR2** | 0.25 | 1.25 | 1.625 |
